# Neural correlates of approach and avoidance tendencies toward physical activity and sedentary stimuli: An fMRI study

**DOI:** 10.1101/2025.01.10.632302

**Authors:** Boris Cheval, Leonardo Ceravolo, Kinga Igloi, David Sander, Myriam Zimmermann, Peter van Ruitenbeek, Matthieu P. Boisgontier

## Abstract

Automatic tendencies toward physical activity and sedentary stimuli are involved in the regulation of physical activity behavior. However, the brain regions underlying these automatic tendencies remain largely unknown. Here, we used an approach-avoidance task and magnetic resonance imaging (MRI) in 42 healthy young adults to investigate whether cortical and subcortical brain regions underpinning reward processing and executive function are associated with these tendencies. At the behavioral level, results showed more errors in avoidance behavior following sedentary stimuli than physical activity stimuli. At the brain level, avoidance behavior following sedentary stimuli was associated with more activation of the motor control network (dorsolateral-prefrontal cortex, primary and secondary motor cortices, somatosensory cortex). In addition, increased activation of the bilateral parahippocampal gyrus — and structural deformation of the right hippocampus - were associated with a tendency toward approaching sedentary stimuli. Together, these results suggest that avoiding sedentary stimuli requires higher levels of behavioral control than avoiding physical activity stimuli.

## Introduction

Exercise is one of the most popular New Year’s resolutions. Unfortunately, this pledge often fails by the month of February (Luciani, 2015), which illustrates the difficulty to engage in physical activity. There is an urgent need to address this inability to be physically active in order to slow the global increase of inactivity (Strain et al., 2024) and achieve the goal of reducing physical inactivity by 15% by 2030 (WHO, 2019). Meanwhile, physical inactivity costs 67.5 billion international dollars each year (Ding et al., 2016) and is responsible for approximately one death every six seconds worldwide (WHO, 2020).

Recent theoretical work has suggested that automatic responses to physical activity and sedentary cues are essential in explaining the gap between intentions to be physically active and actual engagement in physical activity (Brand & Ekkekakis, 2018; Cheval & Boisgontier, 2021; Cheval, Radel, et al., 2018; Cheval et al., 2024; Conroy & Berry, 2017; Maltagliati, Raichlen, et al., 2024). In particular, within the dual-process account of human behavior (Strack & Deutsch, 2004), the Theory of Effort Minimization in Physical Activity (TEMPA) argues that people have an automatic attraction to effort minimization, which may lead individuals to be automatically attracted to sedentary opportunities that arise in their environment (Cheval & Boisgontier, 2021). TEMPA posits that (1) sedentary behaviors are rewarding, and (2) avoiding sedentary behaviors requires more executive control than approaching sedentary behaviors or avoiding physical activity (Cheval & Boisgontier, 2021; Cheval, Radel, et al., 2018).

According to TEMPA’s first postulate, sedentary behavior should be intrinsically rewarding and provide motivational drive to favor that behavior. This drive may be characterized by activation of specific brain regions. However, current neural evidence for the rewarding or motivational value of sedentary behavior is unclear. Some studies support this first postulate (Jackson et al., 2014; Prévost et al., 2010). For example, obese women showed a reduced activation of reward brain areas than lean women when viewing pictures of physical activity, suggesting that higher effort is associated with lower reward (Jackson et al., 2014). In addition, the prospect of energetic expenses was associated with activation in the anterior cingulate cortex and anterior insula, which was interpreted as signaling higher perceived costs (Prévost et al., 2010). However, other studies challenge this first postulate. For example, Crémers et al. (2012) showed that brain areas associated with reward (e.g., insula, pallidum, caudate) and motor control (e.g., dorsolateral prefrontal cortex [DLPFC]) were activated during the mental imagery of brisk walking (compared to lying and standing conditions). Using a go/no-go task toward stimuli depicting physical activity and inactivity, no evidence of activation was shown in brain areas associated with reward processing (Kullmann et al., 2014). Finally, in studies using electroencephalography (EEG), reward-related brain activity showed no evidence supporting that sedentary behavior was rewarding (Cheval, Boisgontier, et al., 2019; Parma et al., 2023). In summary, the neural evidence regarding the rewarding or motivating value of sedentary behavior is inconsistent.

Building on TEMPA’s second postulate, it can be suggested that active avoidance (i.e., moving away from sedentary behavior) requires executive control, involving activation of associated brain areas. In contrast, passive avoidance (i.e., refraining from moving toward sedentary behavior) may specifically depend on inhibitory control. Studies consistently support this second postulate, indirectly validated by large-scale epidemiological studies showing the importance of cognitive function in facilitating and sustaining engagement in physical activity (Cheval, Boisgontier, et al., 2022; Cheval, Orsholits, et al., 2020; Cheval, Rebar, et al., 2019; Cheval et al., 2023; Csajbók et al., 2022; Daly et al., 2015; Sabia et al., 2017). EEG studies provide a more direct support for this postulate (Cheval et al., 2021; Cheval, Daou, et al., 2020; Cheval, Tipura, et al., 2018). For example, avoiding sedentary stimuli, compared to avoiding physical activity stimuli, was associated with larger evoked-related potentials in the frontal cortical areas (Cheval, Tipura, et al., 2018). Similarly, a study using a go/no-go task showed that passively avoiding stimuli representing sedentary behaviors, compared to physical activity, was associated with larger evoked-related potentials in the frontocentral cortex (Cheval et al., 2021; Cheval, Daou, et al., 2020). However, the limited spatial resolution of EEG prevents these studies from precisely identifying the neural networks underlying these automatic responses.

To the best of our knowledge, only one fMRI study has been conducted to investigate brain areas potentially underlying executive control in the processing of physical activity and sedentary stimuli (Kullmann et al., 2014). The results of this study suggest that passively avoiding stimuli related to physical activity is associated with an increased demand on the inhibitory control system (e.g., prefrontal cortex) in patients with anorexia nervosa (Kullmann et al., 2014). However, this association may be explained by the fact that patients with anorexia nervosa often report excessive levels of physical activity (Davis et al., 1997), limiting the generalization of the results to the general population, where a reverse pattern can be expected (Cheval et al., 2021; Cheval, Daou, et al., 2020). Therefore, using MRI to investigate the brain regions underlying executive control in the processing of physical activity and sedentary stimuli in healthy adults is warranted.

### The present study

The aim of the present study was to investigate whether brain regions involved in reward processing and executive control are associated with the processing of stimuli depicting physical activity and sedentary behavior using MRI. Specifically, based on the postulates of TEMPA and previous work, this study focused on brain regions associated with reward processing, such as orbitofrontal cortex, amygdala, and ventral striatum (Corbit & Balleine, 2011; Gottfried et al., 2003; Knutson et al., 2001; Prévost et al., 2012; Roesch & Olson, 2004; Schultz et al., 2000), or with executive control, such as DLPFC, inferior frontal cortex, presupplementary motor area, and basal ganglia (striatum and subthalamic nucleus) (Aron et al., 2007; Aron et al., 2014; Zandbelt & Vink, 2010). To this end, healthy young participants performed an ’implicit’ approach-avoidance task using stimuli depicting avatars running, standing, and sitting during fMRI. In addition, analysis of subcortical structures shapes were associated with the tendency to avoid physical activity of approach sedentary behavior.

### Hypotheses

At the behavioral level, we hypothesized shorter reaction times and/or fewer errors when approaching sedentary stimuli than when approaching physical activity stimuli (HB1). In contrast, we hypothesized longer reaction times and/or more errors when avoiding sedentary stimuli than when avoiding physical activity stimuli (HB2).

At the brain level, we hypothesized increased activity in brain areas associated with reward when approaching compared to avoiding sedentary stimuli (HN1) (contrast: approach sedentary > avoid sedentary). In addition, we hypothesized increased activity in brain areas involved in executive control when avoiding compared to approaching sedentary stimuli (HN2) (contrast: avoid sedentary > approach sedentary) and when avoiding sedentary stimuli compared to avoiding physical activity stimuli (HN3) (contrast: avoid sedentary > avoid physical activity). We also hypothesized that brain activity differences observed in HN3 would not be observed with stimuli depicting very light physical activity (i.e., standing) (contrast: avoid sedentary > avoid neutral) (HN4). Finally, we expected the shape of subcortical brain structures associated with reward processing (i.e. nucleus accumbens, pallidum) and generation of habitual responding (i.e. caudate, putamen) would be associated with the tendency to avoid physical activity and approach sedentary behavior. Other subcortical areas were part of an exploratory analysis.

## Materials and Methods

### Participants

To estimate the sample size required for adequate power (90%) with an alpha level set at 5%, we conducted an a priori power analysis using G*Power 3.1 (Faul et al., 2009). We performed a power analysis for a repeated-measures ANOVA with a small to medium effect size (Cohen’s d = 0.40). We set groups to one, measures to six (action, stimuli), correlations between repeated measures to 0.5, and non-sphericity to one. The power calculation estimated a required *N* of 36, but we aimed to recruit 45 to account for potential data loss due to collection issues.

Exclusion criteria included a history of psychiatric, neurological, or severe mental disorders; use of psychotropic medications, alcohol, or illicit drugs at the time of the study; and any MRI contraindications. In addition, participants were screened to include only those who were right-handed (Oldfield, 1971), could understand French, were >18 years of age, and were free of any medical conditions that would prohibit physical activity without supervision. Smokers were abstinent from tobacco for at least 1.5 hours prior to scanning to reduce the effects of nicotine on the blood oxygen dependent level (BOLD) signal (Jacobsen et al., 2002). Participants read and completed a written informed consent form. The study was approved by the Ethics Committee of the Canton of Geneva, Switzerland (CCER-2019-00065). Participants were compensated with 100 Swiss francs for their participation.

Forty-seven healthy volunteers were recruited. Data from 5 participants were excluded due to the inability to enter the MRI scanner (e.g., presence of piercings, tattoos, or copper intrauterine device). The final sample consisted of 42 participants (31 women, 23.0 ± 3.5 years; body mass index = 21.4 ± 3.0 kg.m-2).

### Experimental paradigm

At least two days prior to the experimental session, participants completed an online questionnaire measuring their laterality (Edinburgh Handedness Inventory) (Oldfield, 1971), usual level of physical activity and sedentary behavior (International Physical Activity Questionnaire) (Craig et al., 2003), motivation for physical activity (i.e., attitudes, intentions, and motivation), exercise dependence (Griffiths et al., 2005), approach-avoidance temperament (Elliot & Thrash, 2010), and demographics (age, sex, height, and weight). Prior to entering the MRI scanner, participants completed a checklist to ensure that they met the requirements to perform a task in the MRI scanner and a questionnaire to assess potential confounding variables (e.g., caffeine, alcohol, and cigarette consumption). An MRI assistant then equipped the participants with the physiological measurements (i.e., respiratory rate, galvanic response, cardiac rhythm) and positioned them in the scanner. Participants were instructed on how to behave during the experiment (e. g., move as little as possible, especially the head). Both foam padding and a strap across the participant’s forehead were used to minimize head movement.

To assess approach-avoidance tendencies and the associated neural activations, participants completed the Visual-Approach/Avoidance-by-the-Self-Task (VAAST) (Rougier et al., 2018) during fMRI. The task was presented using E-Prime (beta 5.0 version) software (Psychology Software Tools Inc.). The MRI sequences included a T1-weighted scan (5 min), a resting state (8 min), the first two functional runs of the VAAST (8 min each), a T2-weighted scan (5 min), the last two functional runs (8 min each), and a reward localizer task (13 min).Finally, participants were paid and debriefed. The entire session lasted approximately 100 minutes.

### Stimuli

Using Unity software, we created stimuli depicting avatars in three distinct postures: active (i.e., running), inactive (i.e., sitting in a cubicle), and an intermediate position (i.e., standing), which will be referred to as ’neutral’ throughout the article. Images were created to match for color, brightness and visual complexity. Specifically, a set of 195 images containing 14 avatars (50% woman) in active, inactive and neutral positions was tested in a pilot study in which 105 participants were asked to rate a random set of 65 pictures. They were asked to rate the extent to which they associated each stimulus with “movement and physically active behavior” (versus “rest and physically inactive behavior”) using two Visual Analogue Scales (VAS1: *Please indicate the extent to which you think this image is associated with a behavior that requires*: 0 = *No physical exertion at all*, 100 = *A lot of physical exertion”*; VAS2:*“Please indicate how closely this image is associated with: 0 = Resting, sedentary behavior, 100 = Moving, very active behavior”*). Participants also rated the credibility (*“How realistic do you think this person’s behavior is? Realistic means that the pictures may resemble to a real-life behavior”*; on a VAS from 0=*behavior not at all realistic*; 100 = *Behavior very realistic*) and the likeability of each picture (“*How likeable/sympathetic do you find the person in this picture? For example, would you like to talk to him/her*”; on a VAS from 0 = *Very unpleasant/antipathetic*, 100 = *Very pleasant/sympathetic*).

The purpose of the pilot study was twofold. First, to ensure that the selected pictures reflected the concept of interest (i.e., movement and physical activity vs rest and physical inactivity). Second, to test whether the selected pictures were equivalent in terms of credibility and pleasantness across categories (i.e., movement versus rest). Based on the results of the pilot study, we selected a total of 84 pictures that included 12 avatars (50% woman) in seven positions (three running positions, three sitting positions, and one standing position). Note that each avatar was represented in the seven positions to ensure a strict equivalence between the conditions (i.e., physically active, physically inactive, and standing).

The selected physical activity-related pictures were evaluated as associated with a significantly higher level of physical effort (72.4 ± 2.52) compared to the sedentary-related pictures (17.45 ± 2.98, *p* < 0.001) and the neutral pictures (38.15 ± 2.01, *p* < 0.001). Similarly, the sedentary-related pictures were evaluated as being associated with a significant lower level of physical effort compared to the neutral pictures (*p* < 0.001). In addition, on average, the pictures were rated as credible (81.48 ± 3.10) and had a moderate level of pleasantness (55.72 ± 7.92). No difference in the level of credibility was observed between physical activity and sedentary pictures (81.63 ± 2.83 and 80.24 ± 2.86 for physical activity and sedentary stimuli, respectively, *p* = 0.089), but neutral pictures were rated as slightly more credible (84.70 ± 2.12) compared to physical activity (*p* = 0.004) or sedentary (*p* < 0.001) pictures. No significant differences in the level of pleasantness were observed between the different types of pictures (56.14 ± 7.53, 55.15 ± 8.16, and 56.20 ± 8.97, for activity, sedentary, and neutral pictures, *p* = 0.850). These results demonstrated the validity of the stimuli in terms of their association with the level of physical effort. It also confirms that that these stimuli were equivalent in terms of pleasantness and credibility, except for the neutral pictures, which were rated slightly more credible than the activity and sedentary pictures.

### The Visual-Approach/Avoidance-by-the-Self-Task (VAAST)

An adapted version of the VAAST was used to measure automatic approach-avoidance tendencies toward physical activity and sedentary behaviors (Rougier et al., 2018). Compared to other approach-avoidance tasks such as the manikin task (Cheval et al., 2015; Cheval et al., 2014; Krieglmeyer & Deutsch, 2010), the VAAST has been shown to produce large and replicable effects. During the task, participants were asked to respond to the format (i.e., portrait vs. landscape format) of the pictures depicting avatars in active (i.e., running position), inactive (i.e., sitting position), and neither active nor inactive (i.e., standing or “neutral” position) positions by pressing the ‘move forward’ or ‘move backward’ buttons three times on an MR-compatible response box (Current Designs Inc., Philadelphia, PA, USA), which was placed beneath the participant’s fingers. Participants were instructed to approach the picture when it appeared in a portrait format, and to avoid it when it appeared in a landscape format (the rule was counterbalanced across participants). Congruent with the participants’ approach or avoidance response, the entire visual environment zoomed in to simulate an approach movement and zoomed out to simulate an avoidance movement. A 30% change after the button press was used to give the impression of walking forward or backward as a consequence of the responses.

The VAAST was administered in four runs. Each run consisted of 54 trials, for a total of 216 trials. Each run included an equal number of trials (i.e., 9) for each of the six conditions representing the interaction between the two main factors Type of action and Type of stimuli (i.e., approach activity, approach neutral, approach sedentary, avoid activity, avoid neutral, and avoid sedentary). The stimuli were pseudorandomized across the runs. To avoid expectancy effects, we varied the duration of the fixation cross (interstimulus interval; 4–8 s) in each trial (Figure 1).

**Figure 1.**
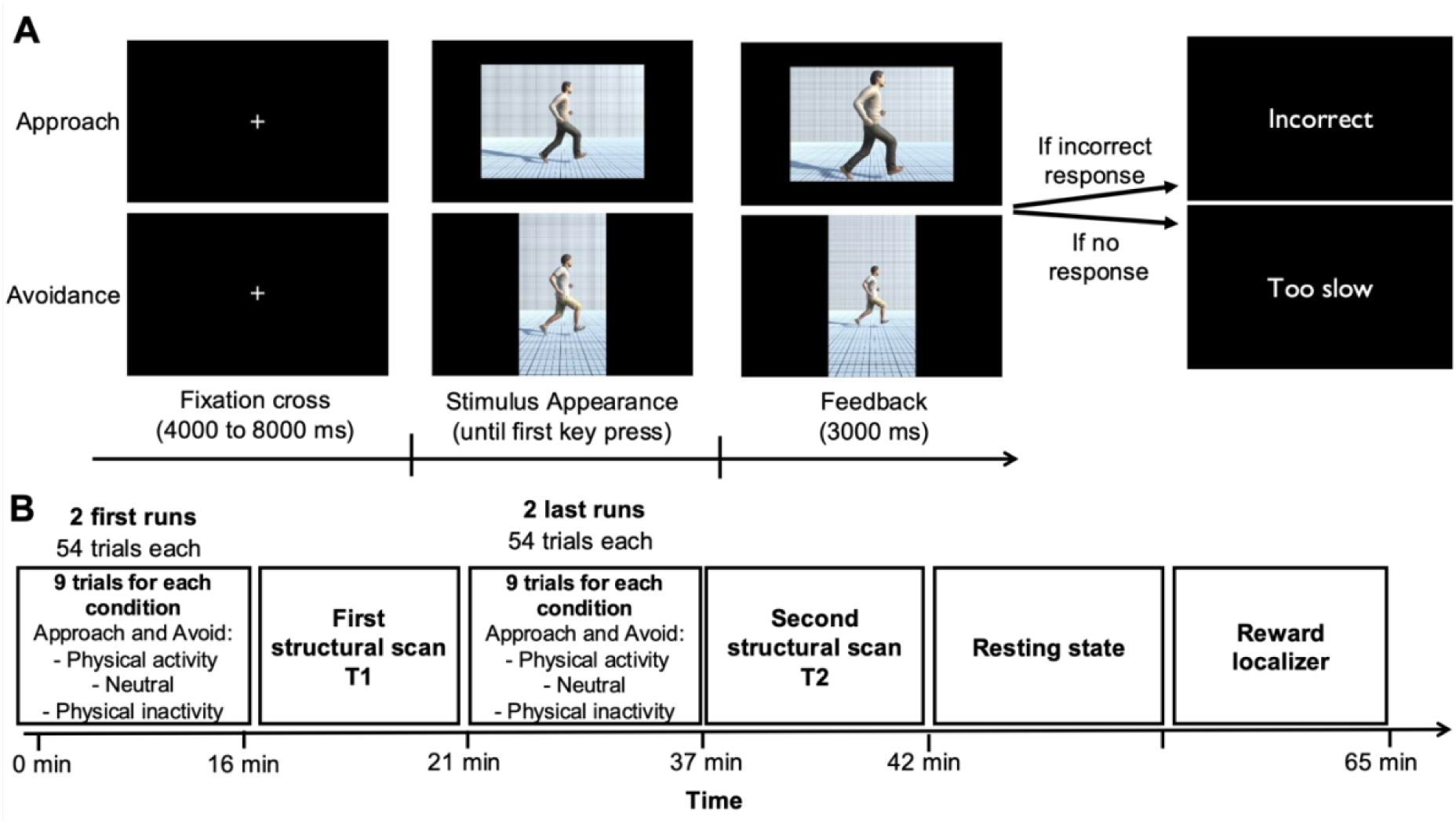
Experimental paradigm. A. the approach-avoidance task. Participants were instructed to quickly approach or avoid pictures depending on their format (i.e., portrait vs landscape format). The six conditions (i.e., approach physical activity, approach neutral, approach physical inactivity, avoid physical activity, avoid neutral, and avoid physical inactivity) were pseudo-randomized across the run. *B. Procedure.* Participants were asked to complete four runs of the approach-avoidance task. Each run was composed of 54 trials, including 9 trials within each of the six conditions.

### Behavioral analyses

Statistical analyses of the behavioral data (i.e., reaction times and errors) were performed using R (R Core Team, 2019). Specifically, mixed-effects models (Baayen et al., 2008; Boisgontier & Cheval, 2016) were used via the lme4 and lmerTest packages (Bates et al., 2014; Kuznetsova et al., 2015) to account for the cross-random structure of the current data (i.e., a random sample of participants crossed with a random sample of stimuli) and thereby correctly estimate the parameters.

To examine participants’ reaction times, the linear mixed-effects models included as fixed factors the type of stimuli (i.e., physical activity, sedentary behaviors, and neutral) and the type of action (i.e., approach, avoidance), and an interaction between these two fixed factors. Participants and stimuli (i.e., pictures) were specified as random factors, and the model also included random effects of the type of action, the type of stimuli, and of their interaction at the participant level. These random parameters allowed the effects of the fixed factors (and of their interaction) on the reaction times to vary across participants. For exploratory analyses, additional models included three-way interactions between usual physical activity level, sedentary craving state, and physical activity craving state with the type of stimulus and the type of action (see Supplementary Material 1 for more details on this measure). These latter models allowed us to examine the extent to which dispositional or situational factors might alter participants’ reaction times to approach (vs. avoid) physical activity, neutral, and sedentary stimuli, as expected by TEMPA (Maltagliati, Fessler, et al., 2024). The same models were applied to errors, except that linear mixed-effects models were replaced by logistic mixed effects models to predict the probability of making an error.

To reduce convergence problems, each model was optimized using the default BOBYQA optimizer (Powell, 2009), the Nelder-Mead optimizer (Nelder & Mead, 1965), the nlimb optimizer from the optimx package (Nash & Varadhan, 2011), and the L-BFGS-B optimizer (see Cheval, Bacelar, et al., 2020; Cheval et al., 2021; Cheval, Maltagliati, et al., 2022; Frossard & Renaud, 2019, for similar procedure). *P* values for the global effect of the factors and of their interaction were reported using likelihood ratio tests comparing models with and without the fixed factors included in the models. Statistical assumptions associated with mixed-effects models (i.e., normality of the residuals, linearity, multicollinearity, and undue influence) were met.

### fMRI data acquisition

Structural and functional imaging was performed at the Brain and Behavior Laboratory (BBL) of the University of Geneva. High-resolution imaging data were acquired on a 3-Tesla whole-body MRI system (Magnetom Tim Trio, Siemens, Erlangen, Germany) equipped with a 12-channel head coil. We used multislice echo planar imaging sequences. For each participant and for each run of the experimental task, 79 functional 2D T2*-weighted echo planar image volumes (EPIs; voxel size = 2.5×2.5×2.5 mm, 48 slices, TR = 600 ms, TE = 32 ms, matrix = 84×84, FoV = 210×210 mm, in-plane resolution = 64×64, FA = 52 degrees) were acquired. Thus, an average of 900 volumes of 48 slices were acquired for each participant. The 192 high-resolution 3D T1-weighted structural images (1mm^3^ isotropic voxels, TR = 1900 ms, TE = 2.27 ms, FA = 9 degrees, FoV = 256×256 mm) were also acquired using a magnetization-prepared rapid acquisition gradient echo sequence.

### fMRI data preprocessing

Functional images were analyzed using Statistical Parametric Mapping software (SPM12, Wellcome Trust Centre for Neuroimaging, London, UK). Preprocessing steps included realignment to the first volume of the time series, normalization to the Montreal Neurological Institute (MNI) space (Collins et al., 1994) and spatial smoothing with an isotropic Gaussian filter of 8 mm full width at half maximum. To remove low-frequency components, we used a high-pass filter with a cutoff frequency of 1/128Hz.

### fMRI data analysis

Data were analyzed using general linear modeling (GLM) as implemented in SPM12 (https://www.fil.ion.ucl.ac.uk/spm/). For the first-level analyses of the experimental task, correctly scored trials of our conditions of interest (design matrix conditions: 1. approach physical activity stimulus; 2. avoid physical activity stimulus; 3. approach sedentary stimulus; 4. avoid sedentary stimulus; 5. approach neutral stimulus; 6. avoid neutral stimulus) and trial-level reaction times were modeled by fitting a boxcar function at the onset of the feedback screen convolved with the canonical hemodynamic response function for 3 sec (duration of the feedback screen). An additional column was added to the design matrix, containing error trials (wrong response trials) and trials for which response times were outside the bounds of percentiles 2 and 98 to remove trials in which participants either pressed too quickly to see the image or did not respond at all. These types of trials were concatenated into a single column per run and only contained on average 2 trials per run. The design matrix therefore included our 6 columns of interest with the corresponding 6 columns of reaction times and the ‘error’ trials and the 6 realignment parameters to account for movement in the data, for a total of 19 columns per run per participant. The four runs were modelled in a single first-level design matrix with runs separated as four different sessions of one participant. Contrasts were computed with the main effect of each of the 6 conditions of interest (value of ‘1’) inversely correlating with reaction times for each condition (value of ‘-1’).

Whole brain group-level statistics were then performed using a 252-lines flexible factorial analysis, in which the first-level simple effects were implemented (42 participants * 6 conditions = 252 files/lines). The model therefore included the factors Participants, Type of action (i.e., approach, avoidance) and Type of stimuli (i.e., physical activity, sedentary behaviors, and neutral). Their interaction was also tested. Independence was set to ‘true’ for the Participants factor and to ‘false’ for the remaining within-factors. Variance estimation was set to ‘unequal’ for all factors because homoscedasticity criteria cannot usually be met for fMRI data (default setting in SPM12). Group-level results of our final contrasts of interest—see hypotheses section – were then corrected for multiple comparisons using a voxel-wise threshold of *p <* .05 with false discovery rate correction (FDR) and an arbitrary cluster extent of k > 10 voxels to remove extremely small clusters of activation. For all analyses, regions were labeled using the latest version of the Automated Anatomical Labelling Atlas (‘AAL3’) (Rolls et al., 2020) and rendered on semi-inflated brains from the CONN toolbox (http://www.nitrc.org/projects/conn).

### Vertex analysis

A partial exploratory analysis was performed to determine presence of an association between the shape of subcortical structures (i.e., nucleus accumbens, amygdala, caudate, hippocampus, pallidum, putamen, and thalamus) and a behavioral bias towards approaching sedentary behavior, and between the shape of these structures and a behavioral bias towards avoiding physical activity. Tendency towards sedentary behavior is represented by the difference between responses (speed and accuracy) representing approaching sedentary behavior and avoiding sedentary behavior (i.e. sedentary approach – sedentary avoidance). Tendency towards avoiding physical activity is represented by the difference between responses (speed and accuracy) representing avoiding physical activity and avoiding sedentary behavior (i.e. activity avoidance – sedentary avoidance). For reaction times the signed was flipped to have higher positive scores represent larger bias.

The individual structure’s shape is represented by a mesh consisting of vertices. These are compared an average mesh to calculate inward and outward deformations of the individual structure. To obtain these measures, first T1 weighted images were reoriented to standard orientation. Next, structures were segmented from the T1 weighted images using FMRIB’s Integrated Registration Segmentation Toolkit (FSL FIRST; Patenaude et al., 2011) in FSL version 6.0.7.13 (Jenkinson et al., 2012; Smith et al., 2004; Woolrich et al., 2009). As a sub-step, T1 images are registered to normalized space. Accuracy of the registrations were visually inspected for all participants in using the ‘slicesdir’ command to create coronal, sagittal, and horizontal slices. Subsequently, vertex analysis (FSL) was used to indicate the exact location of the relation between subregional grey matter structure and behavioral tendencies. The vertices represent the signed, perpendicular distance from the average surface. Negative and positive values reflect inward (i.e., local atrophy) and outward (i.e., local expansion) deformation of the structures, respectively. FSL FIRST vertex analysis (Patenaude et al., 2011) restricts topology of the structures and preserves inter-participant vertex correspondence, enabling a vertex-wise comparison of differences between conditions in the association with behavioral tendencies. The regression models using behavior tendencies predicting structural deviations from the mesh representing average shape were created and tested for significance using permutation-based non-parametric tests (FSL randomise, 10,000 draws, p < 0.05, TFCE applied, FWE corrected) (Smith & Nichols, 2009).

## Results

### Descriptive results

Table 1 shows the characteristics of the participants and reports the reaction times to approach and avoid stimuli depicting physical activity, neutral, and sedentary stimuli, as well as the approach bias scores (i.e., reaction times to avoid - reaction times to approach) for each type of stimulus. On average, reaction times within each condition were < 700 ms, and strongly correlated with each other (Pearson’s Rs between .83 and .95, *ps* < .001). Error rates were on average about 5% (± 6%) for avoiding physical activity, 6% (± 9%) for approaching neutral stimuli, and about 7% for the other conditions (standard deviations ranged from 6% to 9%).

**Table 1.** Descriptive statistics.

| N = 42 | Mean | SD |
| --- | --- | --- |
| Age (years) | 23.0 | 3.5 |
| Gender (number; %) |  |  |
| Women | 31 | 74% |
| Men | 11 | 26% |
| Body Mass Index | 21.4 | 3.0 |
| Craving for sedentary behaviors | 4.3 | 1.6 |
| Craving for physical activity behaviors | 3.7 | 1.7 |
| Usual level of physical activity (min per week) | 285.9 | 293.1 |
| **Reaction times (ms)** |  |  |
| Approach physical activity | 666.5 | 111.4 |
| Approach neutral | 668.7 | 117.5 |
| Approach sedentary behaviors | 677.4 | 116.5 |
| Avoid physical activity | 659.6 | 102.8 |
| Avoid neutral | 659.4 | 107.6 |
| Avoid sedentary behaviors | 669.7 | 114.3 |
| **Approach biases (ms)** |  |  |
| Approach bias toward physical activity | -6.9 | 47.2 |
| Approach bias toward neutral stimuli | -9.3 | 65.6 |
| Approach bias toward sedentary behaviors | -7.7 | 65.7 |
| **Errors** |  |  |
| Approach physical activity | 7% | 8% |
| Approach neutral | 6% | 9% |
| Approach sedentary behaviors | 7% | 7% |
| Avoid physical activity | 5% | 6% |
| Avoid neutral | 7% | 8% |
| Avoid sedentary behaviors | 7% | 8% |

*Notes.* SD = standard deviation; ms = milliseconds ; min = minutes

### Reaction Times and Error Rates in the Approach-Avoidance Task

#### Reaction times

The results of the linear mixed-effects models showed no main effect either of stimulus type (*p-*value for global effect = 0.164) or action type (*p*-value for global effect = 0.160). Also, the two-way interaction between stimulus type and action type was also not significant (*p*-value for global effect = 0.965). Simple effects tests further confirmed that reaction times to approach (vs. avoid) physically active stimuli were not statistically different from reaction times to approach (vs. avoid) sedentary stimuli (*p* = .851) (Table 2). Similarly, reaction times to approach (vs. avoid) neutral stimuli were not statistically different from the reaction times to approach (vs. avoid) sedentary stimuli (*p* = .802) or physically active stimuli (*p* = .661).

**Table 2.** Results of the linear mixed-effects models predicting the reaction times as a function of action type (approach vs. avoidance) and stimulus type (physical activity vs. neutral vs. sedentary).

| **N = 40\*** | **b (CI)** | **p** |
| --- | --- | --- |
| **Fixed Effects** |  |  |
| Intercept | **666.1 (627.9;704)** | **<.001** |
| **Stimuli (ref. physical activity)** |  |  |
| Neutral | 2.7 (-11.7;17.2) | .711 |
| Sedentary | 7.9 (-6.0;21.9) | .267 |
| **Action (ref. approach)** |  |  |
| Avoidance | -7.0 (-23.6;9.6) | .410 |
| **Stimuli (ref. physical activity) x Action (ref. approach)** |  |  |
| Avoidance x neutral | -4.1 (-11.7;14.2) | .661 |
| Avoidance x Sedentary | -1.8 (-22.4;16.6) | .851 |
| **Covariates** |  |  |
| Age | -12.3 (-46.4;21.7) | .483 |
| Sex | -5.1 (-78.3;68.1) | .892 |
| BMI | 1.3 (-33.0;35.6) | .940 |
| **Random Effects** |  |  |
| **Participants** |  |  |
| Intercept | 10710.07 |  |
| Stimuli sedentary | 54.48 |  |
| Stimuli neutral | 8.24 |  |
| Action Avoid | 1109.29 |  |
| Corr. (Intercept, stimuli sedentary) | 0.07 |  |
| Corr. (Intercept, stimuli neutral) | 0.97 |  |
| Corr. (Intercept, action avoidance) | -0.30 |  |
| Corr. (Stimuli sedentary; stimuli neutral) | -0.16 |  |
| Corr. (Stimuli sedentary; action avoidance) | 0.93 |  |
| Corr. (Stimuli neutral; action avoidance) | -0.51 |  |
| **Stimuli** |  |  |
| Intercept | 94.87 |  |
| Residual | 28551.56 |  |
| R<sup>2</sup> | Conditional .005 |  |
|  | Marginal =.275 |  |

*Notes.* 95CI = confidence intervals at 95%. \*Two participants were not included in the analyses because they were an issue regarding the recording of their behavioral data.

**Table 3.** Results of the logistic mixed-effects models predicting the risk of error in the approach-avoidance task as a function of action type (approach vs. avoidance) and stimuli type (physical activity vs. neutral vs. sedentary).

| **N = 40** | **OR (CI)** | **p** |
| --- | --- | --- |
| **Fixed Effects** |  |  |
| Intercept | **0.06 (0.04;0.08)** | **<.001** |
| **Stimuli (ref. physical activity)** |  |  |
| Neutral | 0.80 (0.59;1.08) | .149 |
| Physical inactivity | 0.85 (0.63;1.16) | .308 |
| **Action (ref. approach)** |  |  |
| Avoidance | 0.73 (0.53;1.03) | .071 |
| **Stimuli (ref. physical activity) x Action (ref. approach)** |  |  |
| Avoidance x neutral | **1.57 (1.01;2.43)** | **.044** |
| Avoidance x Physical inactivity | **1.64 (1.06;2.54)** | **.025** |
| **Covariates** |  |  |
| Age | 1.04 (0.77;1.40) | .798 |
| Sex | 0.94 (0.49;1.81) | .852 |
| BMI | 0.90 (0.67;1.23) | .516 |
| **Random Effects** |  |  |
| **Participants** |  |  |
| Intercept | 0.68 |  |
| Stimuli physical inactivity | 0.01 |  |
| Stimuli neutral | 0.02 |  |
| Action Avoid | 0.01 |  |
| Corr. (Intercept, stimuli physical inactivity) | 0.65 |  |
| Corr. (Intercept, stimuli neutral) | 1.00 |  |
| Corr. (Intercept, action avoidance) | -0.37 |  |
| Corr. (Stimuli physical inactivity; stimuli neutral) | 0.57 |  |
| Corr. (Stimuli physical inactivity; action avoidance) | -0.95 |  |
| Corr. (Stimuli neutral; action avoidance) | -0.28 |  |
| **Stimuli** |  |  |
| Intercept | null |  |
| R<sup>2</sup> | Conditional .006 |  |
|  | Marginal =.192 |  |

Notes. OR= odds ratio; 95CI = confidence intervals at 95%. Note that the models estimated a null variance for the random intercept of the stimuli. The models with or without this parameter lead to consistent results. \*Two participants were not included in the analyses because they were an issue regarding the recording of their behavioral data.

#### Errors

The results of the logistic mixed-effects models showed no main effect either of stimulus type (*p-*value for global effect = 0.784) or action type (*p*-value for global effect = 0.995). However, although the main effect of the interaction between stimulus type and action type was only marginal (*p*-value for global effect = 0.091), the results showed that the probability of error when avoiding (vs. approaching) physical activity stimuli was statistically different from the probability of error when avoiding (vs. approaching) sedentary stimuli (OR = 1.64, 95%CI = 1.06 – 2.54, *p* = .025) – participants made more errors when instructed to avoid stimuli depicting sedentary behaviors than when instructed to avoid stimuli depicting physical activity. No difference was observed in the approach condition (Figure 2). The same pattern of effect was found between neutral and physical activity stimuli (OR = 1.57, 95%CI = 1.01 – 2.43, *p* = .044).

**Figure 2.**
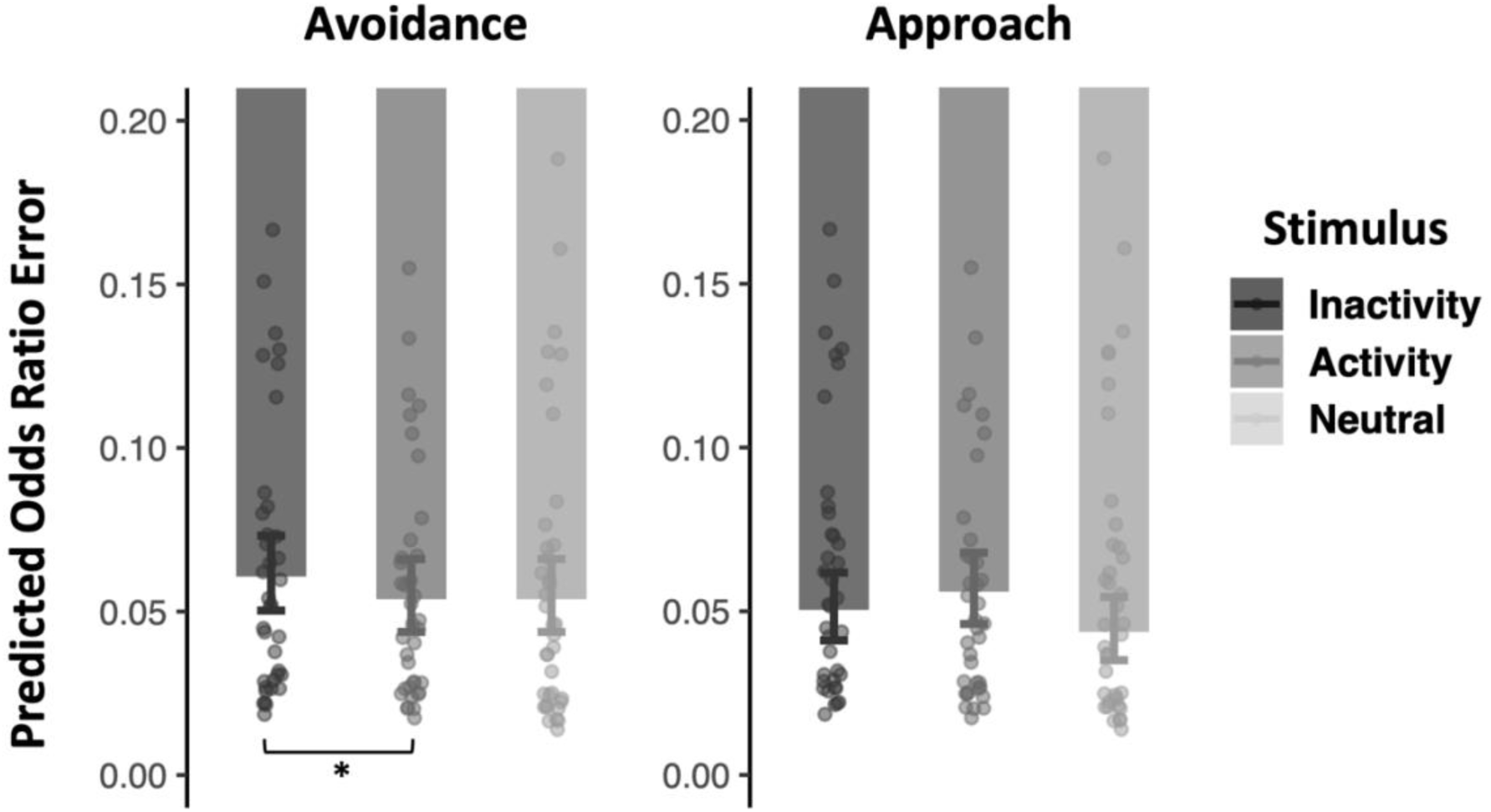
Results of the logistic mixed-effects models. Predicted odds ratio of a failure to avoid or approach stimuli depicting physical activity, neutral, and sedentary behaviors. Dots represent mean response times for each participant as a function of stimulus type (i.e., physical activity vs. sedentary behaviors vs. neutral stimuli). Error bars represent the standard errors around the mean.

### Physical activity engagement and craving for physical activity

#### Reaction times

Results did not show that usual physical activity engagement or craving for physical activity significantly moderated the effect of action, stimulus type, or the interaction between these two factors (see Supplementary Material 2). However, the results showed that reaction time differences between responses following physical activity and sedentary stimuli were moderated by the state of craving for sedentary behaviors (b = −22.0, 95%CI = −35.0 – −9.0, *p* < .001) – participants responded faster to sedentary than to physical activity stimuli when their craving for sedentary behaviors was high, but were slower when the craving for sedentary behaviors was low.

#### Error

Results did not show that usual physical activity, craving for physical activity, or craving for sedentary behaviors significantly moderated the effect of action, stimulus type, or the interaction between these two factors (see Supplementary Material 3).

### Neural Activity Associated with the Avoidance of Sedentary Stimuli

#### Approach sedentary > Avoid Sedentary (HN1)

More activity was observed in the left posterior middle temporal gyrus (Figure 3A), bilateral parahippocampal gyrus (Figure 3DEF), primary and secondary visual cortex (Figure 3BCDFH) when participants approached compared to avoid sedentary stimuli.

**Figure 3.**
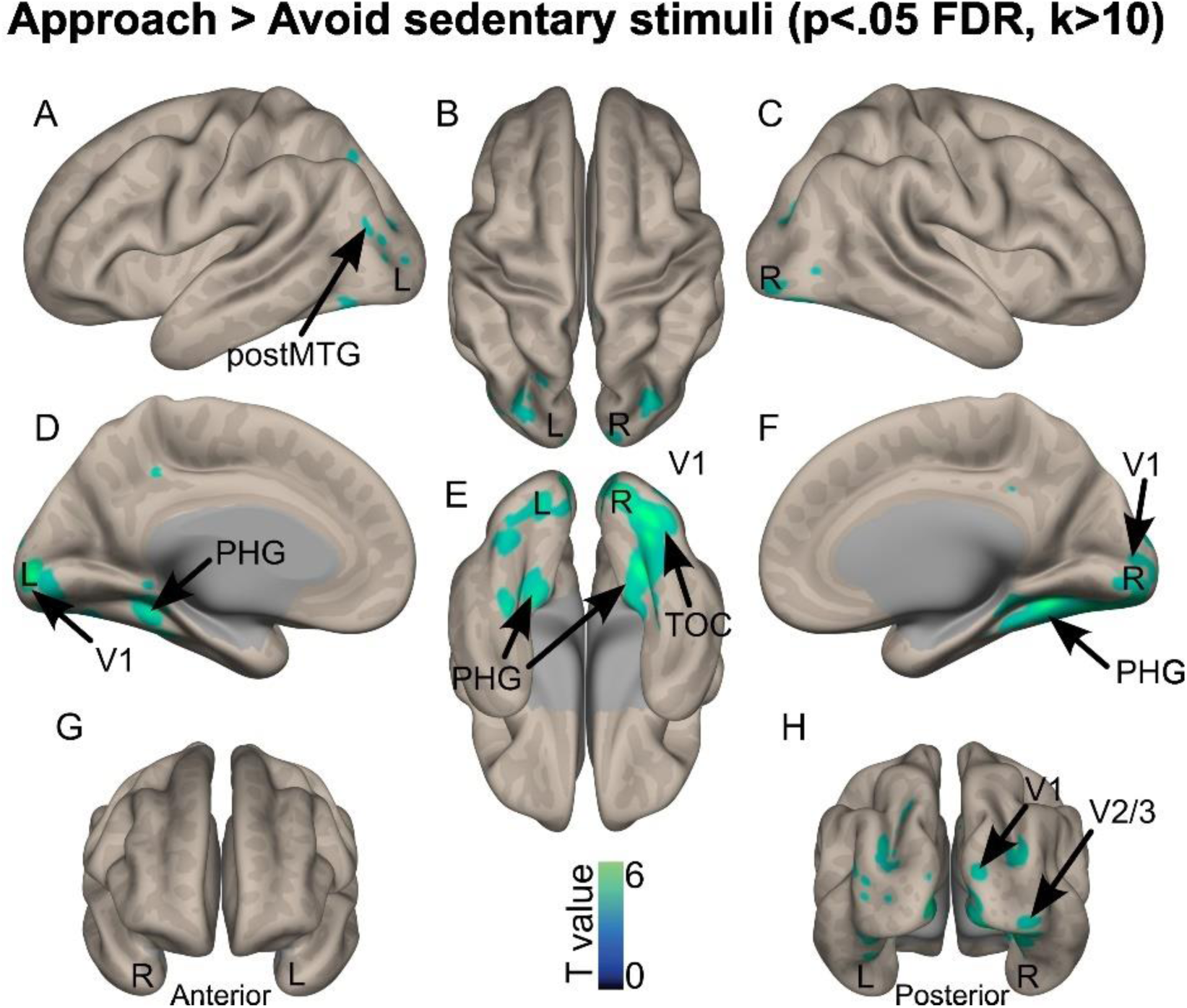
Brain activations when approaching vs. avoiding sedentary stimuli, corrected for multiple comparisons (whole-brain voxel-wise *p*<.05 FDR, k>10 voxels). The color bar represents the statistical T value. V1: primary visual cortex; V2/3: secondary visual cortex; postMTG: posterior part of the middle temporal gyrus; TOC: temporo-occipital cortex; PHG: parahippocampal gyrus. L: left hemisphere; R: right hemisphere.

#### Avoid Sedentary > Approach Sedentary (HN2)

More activity was observed in a widespread network of bilateral brain areas, including the primary motor cortex (Figure 4ABC), the supplementary motor area (Figure 4DF), the primary somatosensory cortex and the bilateral dorsolateral prefrontal cortex (Figure 4ABCG), the bilateral insula (Figure 4AC), the inferior frontal gyrus *pars triangularis* (Figure 4C) and the putamen (Figure 4E), when participants avoided sedentary stimuli as compared to when participants approached sedentary stimuli.

**Figure 4.**
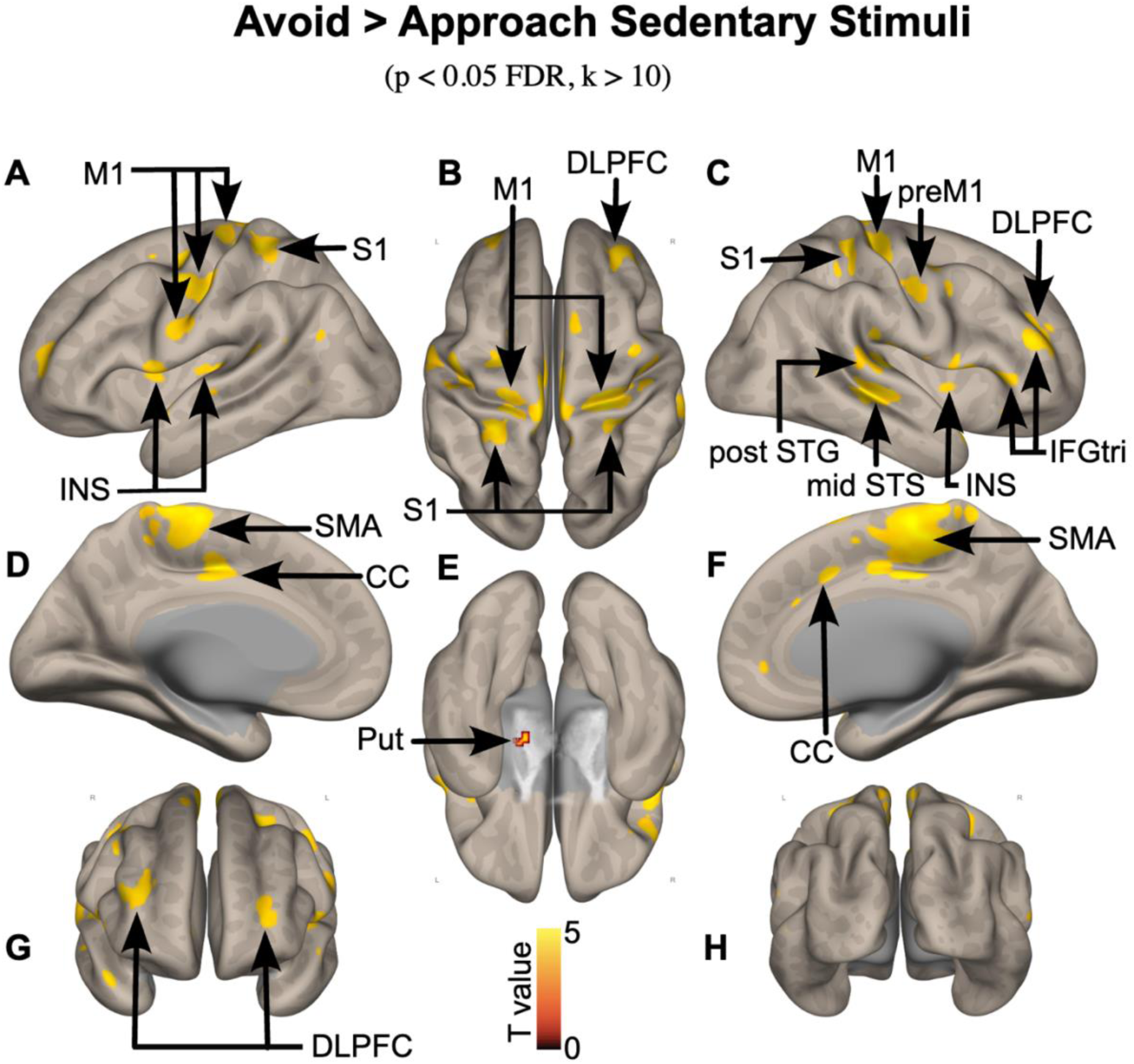
Brain activations when avoiding vs. approaching sedentary stimuli, corrected for multiple comparisons (whole-brain voxel-wise *p*<.05 FDR, k>10 voxels). The color bar represents the statistical T value. postSTG: posterior part of the superior temporal gyrus; midSTS: mid part of the superior temporal sulcus; M1: primary motor cortex; S1: primary somatosensory cortex; INS: insula; DLPFC: dorsolateral prefrontal cortex; IFGtri: inferior frontal gyrus *pars triangularis*; SMA: supplementary motor area; CC: cingulate cortex. L: left hemisphere; R: right hemisphere.

#### Avoid Sedentary > Avoid Physical Activity (HN3)

More activity was observed in the left primary motor cortex, insula, anterior superior temporal sulcus (STS), posterior middle temporal gyrus (MTG; Figure 5AB), right posterior MTG, superior temporal gyrus, mid STS, posterior cingulate cortex, and dorsolateral prefrontal cortex (MTG; Figure 5CFG). Subcortical activations were also observed especially in the bilateral putamen and in the left thalamus (MTG; Figure 5E), when participants avoided sedentary behavior than when the avoided physical activity stimuli.

**Figure 5.**
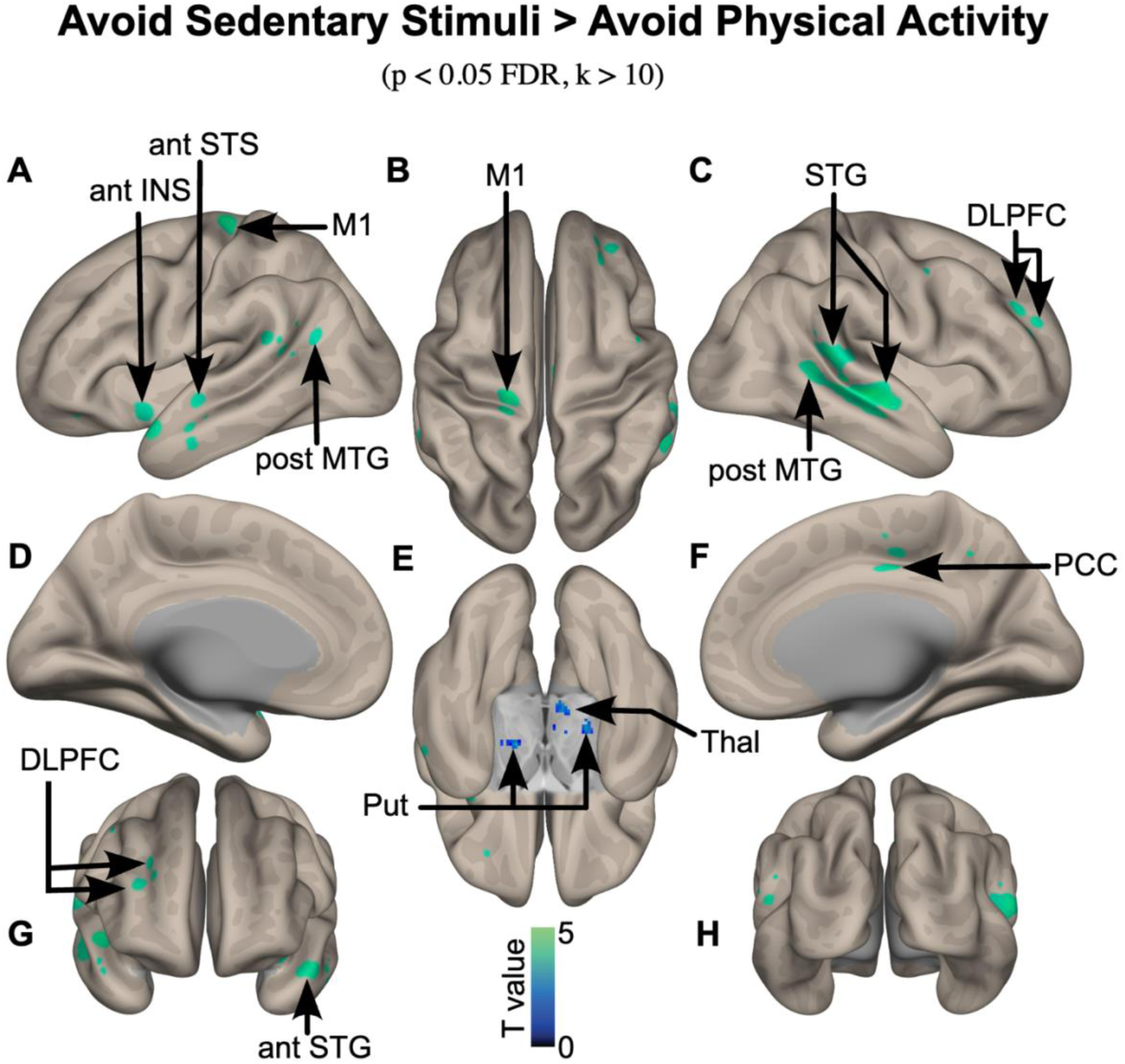
Brain activations when avoiding sedentary stimuli vs. approaching physical activity stimuli, corrected for multiple comparisons (whole-brain voxel-wise *p*<.05 FDR, k>10 voxels). The color bar represents the statistical T value. postSTG: posterior part of the superior temporal gyrus; postMTG: posterior part of the middle temporal gyrus; midSTS: mid part of the superior temporal sulcus; antSTS: anterior part of the superior temporal sulcus; midSTG: mid part of the superior temporal gyrus; M1: primary motor cortex; antINS: insula, anterior part; DLPFC: dorsolateral prefrontal cortex; PCC: posterior cingulate cortex; Thal: thalamus; Put: putamen. L: left hemisphere; R: right hemisphere.

#### Avoid Sedentary > Avoid Neutral (HN4)

More activation was found in the left primary visual cortex, associative visual cortex, temporo-occipital cortex and superior parietal lobule as well as in the right hemisphere in the similar regions, when participants avoided sedentary stimuli as compared to when participants avoided light physical activity stimuli (See Supplementary Materials 4).

See also Supplementary Materials 5 2 for detailed coordinates of the clusters presented in this section.

### Associations Between Subcortical Structure Shapes and Behavioral Bias

The association between subcortical structures’ shape and sedentary behavior tendency was assessed by error and reaction time measures predicting the size of the deformations of that shape. Greater tendency towards sedentary behavior as assessed with reaction time was predictive for larger outward deformations of the right ventral hippocampus (Figure 6). The bias as assessed using error data did not show such significant association. In addition, no other subcortical structure was significantly associated with the sedentary behavior bias. No significant association was observed between a behavioral bias towards avoiding physical activity and the shape of the assessed subcortical structures.

**Figure 6.**
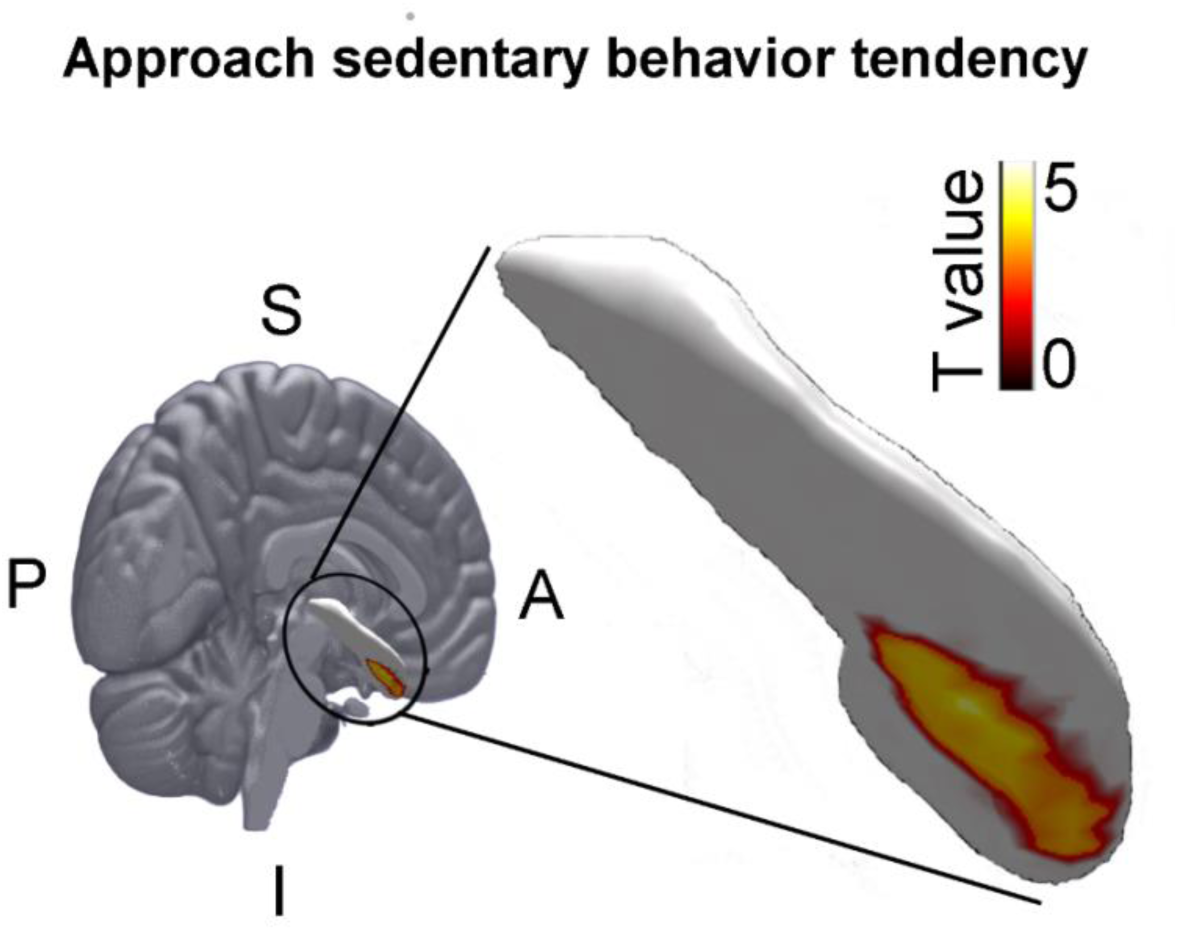
Significant (P < .05, FWE corrected) positive association between deformation of the right hippocampus and the behavioral tendency towards sedentary behavior (average reaction time for approaching sedentary behavior trial < reaction time for avoiding sedentary behavior trials). The extent to which approaching sedentary behavior is easier relative to avoiding it, is associated with an outward deformation of the inferior/anterior right hippocampus. S: Superior, I: Inferior, P: Posterior, A: Anterior

## Discussion

### Main Findings

This study used an approach-avoidance task during fMRI and provides evidence that avoiding sedentary stimuli requires higher levels of behavioral control than avoiding physical activity stimuli. In addition, the outward deformation of the right ventral/anterior hippocampus was associated with a behavioral tendency towards sedentary behavior. These neural results are consistent with behavioral data showing that participants made more errors when avoiding sedentary stimuli than when avoiding physical activity stimuli. Hence, these findings are consistent with the TEMPA’s postulate that avoiding sedentary behaviors requires more executive control than approaching sedentary behaviors or avoiding physical activity, while they did not provide support for the postulate regarding the rewarding value of sedentary behaviors.

### Comparison With Other Studies

#### Behavioral results

Participants made more errors when avoiding sedentary stimuli than when avoiding physical activity stimuli (HB2). This finding is consistent with previous literature that has shown, using a go/no-go task (Duckworth & Kern, 2011), that participants made more commission errors (i.e., a failure to refrain from responding to a “no-go” stimulus) when responding to sedentary stimuli compared to physical activity stimuli (Cheval, Daou, et al., 2020). Thus, these behavioral data provide support for the suggestion that more executive control is required for the avoidance than approach of sedentary opportunities.

However, our results showed no significant effects of stimulus type, action type, or their interaction on participants’ reaction times. This finding contrasts with previous literature that has repetitively shown that participants are faster when approaching compared to avoiding physical activity stimuli, whereas they are faster when avoiding compared to approaching physical inactivity stimuli (Cheval et al., 2015; Cheval et al., 2014; Cheval, Tipura, et al., 2018; Farajzadeh et al., 2023; Hannan et al., 2019; Moffitt et al., 2019). This discrepancy can be explained by the specificity of the task we used in the current study. Specifically, previous studies relied on an explicit approach-avoidance task in which participants were instructed to respond to the content of the picture – to approach or avoid depending on the stimulus type (physical activity or sedentary behavior). In contrast, here we used an ‘implicit’ approach-avoidance task in which participants were instructed to respond to the format of the pictures – to approach or avoid depending on whether the picture appeared in portrait vs. landscape format. A review of the literature found that the implicit stimulus evaluation typically produces smaller effects than explicit stimulus evaluation (Phaf et al., 2014). Accordingly, the reliance on implicit instructions may largely explain why the usual approach tendency toward physical activity and avoidance tendency toward sedentary behavior were not found.

Exploratory analyses further revealed that the state of craving for sedentary behaviors significantly moderated participants’ reaction times in the task. Specifically, greater craving for sedentary behaviors reduced the reaction times in response to sedentary stimuli relative to physical activity stimuli, regardless of the type of action required (i.e., approach or avoidance). These shorter reaction times may be explained by the fact that participants in a state of craving for sedentary behaviors may be more attentive to stimuli associated with such behaviors. This finding is consistent with previous studies showing that attention is biased toward stimuli that are particularly relevant to participant’s current concerns (Cheval, Miller, et al., 2020; Pool et al., 2016). Accordingly, these findings may suggest that physical inactivity stimuli may be particularly relevant to the concerns of individuals who self-report a desire to engage in sedentary behaviors.

#### Neural results

fMRI results showed more activation of a motor control network including primary motor cortex, supplementary motor area, primary somatosensory cortex and dorsolateral prefrontal cortex when participants avoided sedentary stimuli as compared to when participants approached sedentary stimuli. This result suggests that avoiding sedentary behavior requires to deliberately plan and implement the motor action, taking more effort, compared to approach sedentary behavior. However, it is important to note that while this effect was observed specifically for sedentary stimulus contrasts, the conjunction analysis showed no significant differences when comparing sedentary, neutral, and PA stimuli. This calls for caution regarding the specificity of the effect observed for sedentary stimuli. That said, this suggestion is supported by the larger activation observed in the posterior cingulate cortex and DLPFC when participants avoided sedentary behavior compared to when the avoided physical activity stimuli, which could be related to higher resources required for conflict monitoring as well as action planning and implementation. These observations are consistent with previous EEG studies that have shown, using either an approach-avoidance task (Krieglmeyer & Deutsch, 2010) or a go/no-go task (Duckworth & Kern, 2011), that “not going to” or “avoiding” a sedentary stimulus requires greater behavioral control than “not going to” or “avoiding” a physical activity stimulus, as indicated by larger evoked-related potentials in the medial frontal cortex and frontocentral cortex (Cheval et al., 2021; Cheval, Tipura, et al., 2018).

The observed positive association between the outward deformation of the right hippocampus and the tendency to approach sedentary behavior was unexpected, as this structure was a-priori not associated with motivation or reward-based information processing. To potentially explain these findings, it can be argued that the judgement of stimuli being presented in a portrait or landscape format may have been a confounding factor. The currently perceived function of the hippocampus is to encode spatial and temporal contexts of episodes, constructing a cognitive map (Epstein et al., 2017). In particular, the right hippocampus has been shown to be involved in spatial task performance (Klur et al., 2009). In support of our findings, Hernández et al. (2017) performed an analysis similar to the one presented here, linking cognitive function to hippocampal deformation. They observed that a similar subregion of the right hippocampus was specifically associated with spatial memory performance. To test whether judging the stimulus orientation and/or associated movement acted as a confounder, we performed an additional analysis in which we assessed the association between the reaction time difference between approach vs. avoidance of neutral images and the structural deformation of the right hippocampus. Such association between hippocampal structure and reaction time in the neutral condition, may indicate that the observed effect is driven by the orientation of the stimulus and/or associated movement. This analysis did not show any significant association, providing no evidence that the judgement of the spatial orientation was driving the effect. An alternative explanation may be that the currently observed associations reflect an emotion-based decision to engage in approach or avoidance behavior. The presently observed location of the association with sedentary behavior tendency is mostly ventral/anterior, and this subregion of the hippocampus is associated with the processing of stress, emotion and affect. Therefore, speculatively, a larger hippocampal capacity to process intrinsically rewarding events may lead to faster responses such as those observed here.

### Limitations and Strengths

This study has several limitations to consider. First, the experimental setup required participants to lie down, which may have influenced their evaluation of the stimuli and reduced ecological validity. Second, the study’s correlational design, without experimental manipulation, limits the ability to establish causal relationships. Third, the use of self-reported measures to assess usual physical activity introduces potential biases and may partially explain the absence of a moderating effect. Fourth, while the stimuli were validated and relevant to the concept of effort, they cannot fully capture the complexity of effort-related behaviors in real-world contexts. Despite these limitations, the study has notable strengths. The use of fMRI allowed for precise identification of spatial patterns of brain activity, providing valuable insights into the neural mechanisms underlying TEMPA. The design incorporated numerous repetitions within each condition and used a high temporal resolution, optimizing the reliability and quality of the data. The validated stimuli directly addressed the concept of effort, enhancing the study’s relevance, and the well-validated approach-avoidance task added methodological rigor.

## Conclusion

This study provides new insights into the neural mechanisms underlying the difficulty to avoid sedentary stimuli. Behavioral results showed that participants made more errors when avoiding sedentary stimuli compared to physical activity stimuli. Neural results showed greater activation observed in brain regions associated with motor control, conflict monitoring, and action planning when avoiding sedentary stimuli. Altogether, these results suggest that executive control plays an important role in overcoming an inclination toward low-effort behaviors, as proposed by TEMPA.

## Declarations

### Ethical approval

This study was approved by the Ethics Committee of Geneva Canton, Switzerland (CCER-2019-00065).

### Consent to participate

All the participants agreed to participate and signed a written informed consent.

### Consent for publication

All the authors have agreed to the by-line order and to the submission of the manuscript in this form.

### Contributors

BC, MPB designed the study. Data were collected by Master students under the supervision of BC and DS. BC, LC, PvR analyzed the data. BC, KI, DS, PvR, MPB drafted the manuscript. All authors critically appraised and approved the final version of the manuscript.

## Acknowledgments

We extend our gratitude to Fares Alouf and Michael Cecconi for their support during data collection.

## References

Aron, A. R., Durston, S., Eagle, D. M., Logan, G. D., Stinear, C. M., & Stuphorn, V. (2007). Converging evidence for a fronto-basal-ganglia network for inhibitory control of action and cognition. Journal of Neuroscience, 27(44), 11860–11864. 10.1523/JNEUROSCI.3644-07.2007

Aron, A. R., Robbins, T. W., & Poldrack, R. A. (2014). Inhibition and the right inferior frontal cortex: one decade on. Trends in Cognitive Sciences, 18(4), 177–185. 10.1016/j.tics.2013.12.003

Baayen, R. H., Davidson, D. J., & Bates, D. M. (2008). Mixed-effects modeling with crossed random effects for subjects and items. Journal of Memory and Language, 59(4), 390–412. 10.1016/j.jml.2007.12.005

Bates, D., Mächler, M., Bolker, B., & Walker, S. (2014). Fitting linear mixed-effects models using lme4. Journal of statistical software, 67(1). 10.18637/jss.v067.i01

Boisgontier, M. P., & Cheval, B. (2016). The anova to mixed model transition. Neuroscience & Biobehavioral Reviews, 68, 1004–1005.

Brand, R., & Ekkekakis, P. (2018). Affective–Reflective Theory of physical inactivity and exercise. German Journal of Exercise and Sport Research, 48(1), 48–58. 10.1007/s12662-017-0477-9

Cheval, B., Bacelar, M., Daou, M., Cabral, A., Parma, J., Forestier, C., Orsholits, D., Sander, D., Boisgontier, M., & Miller, M. W., (2020). Higher inhibitory control is required to escape the innate attraction to effort minimization. Psychology of Sport and Exercise, 51, 101781.

Cheval, B., Boisgontier, M., Sieber, S., Ihle, A., Orsholits, D., Forestier, C., Sander, D., & Chalabaev, A. (2022). Cognitive functions and physical activity in aging when energy is lacking. European Journal of Ageing, 19, 533–544. 10.1007/s10433-021-00654-2

Cheval, B., & Boisgontier, M. P. (2021). The theory of effort minimization in physical activity. Exercise and Sport Sciences Reviews, 49(3), 168–178. 10.1249/JES.0000000000000252

Cheval, B., Boisgontier, M. P., Bacelar, M. F., Feiss, R., & Miller, M. W. (2019). Opportunities to sit and stand trigger equivalent reward-related brain activity. International Journal of Psychophysiology, 141, 9–17. 10.1016/j.ijpsycho.2019.04.009

Cheval, B., Cabral, D. A. R., Daou, M., Bacelar, M., Parma, J. O., Forestier, C., Orsholits, D., Maltagliati, S., Sander, D., & Boisgontier, M. P. (2021). Inhibitory control elicited by physical activity and inactivity stimuli: An EEG study. Motivation Science, 7(4), 386–389. 10.1037/mot0000236

Cheval, B., Daou, M., Cabral, D. A. R., Bacelar, M., Parma, J. O., Forestier, C., Orsholits, D., Sander, D., Boisgontier, M. P., & Miller, M. W. (2020). Higher inhibitory control is required to escape the innate attraction to effort minimization. Psychology of Sport and Exercise, 51, 101781. 10.1016/j.psychsport.2020.101781

Cheval, B., Maltagliati, S., Fessler, L., Farajzadeh, A., Ben Abdallah, S., Vogt, F., Dubessy, M., Lacour, M., Miller, M. W., Sander, D., & Boisgontier, M. P. (2022). Physical effort biases the perceived pleasantness of neutral faces: A virtual reality study. Psychology of Sport and Exercise.

Cheval, B., Miller, M. W., Orsholits, D., Berry, T., Sander, D., & Boisgontier, M. P. (2020). Physically active individuals look for more: an eye-tracking study of attentional bias. Psychophysiology, 57(6), e13582. 10.1111/psyp.13582

Cheval, B., Orsholits, D., Sieber, S., Courvoisier, D. C., Cullati, S., & Boisgontier, M. P. (2020). Relationship between decline in cognitive resources and physical activity. Health Psychology, 39(6), 519–528. 10.1037/hea0000857

Cheval, B., Radel, R., Neva, J. L., Boyd, L. A., Swinnen, S. P., Sander, D., & Boisgontier, M. P. (2018). Behavioral and neural evidence of the rewarding value of exercise behaviors: a systematic review. Sports Medicine, 48(6), 1389–1404. 10.1007/s40279-018-0898-0

Cheval, B., Rebar, A. L., Miller, M. M., Sieber, S., Orsholits, D., Baranyi, G., Courvoisier, D. C., Cullati, S., Sander, D., & Boisgontier, M. P. (2019). Cognitive resources moderate the adverse impact of poor neighborhood conditions on physical activity. Preventive Medicine, 126, 105741. 10.1016/j.ypmed.2019.05.029

Cheval, B., Saoudi, I., Maltagliati, S., Fessler, L., Farajzadeh, A., Sieber, S., Cullati, S., & Boisgontier, M. (2023). Initial status and change in cognitive function mediate the association between academic education and physical activity in adults over 50 years of age. Psychology and aging, 38(6), 494–507. 10.1037/pag0000749

Cheval, B., Sarrazin, P., Isoard-Gautheur, S., Radel, R., & Friese, M. (2015). Reflective and impulsive processes explain (in)effectiveness of messages promoting physical activity: a randomized controlled trial. Health Psychology, 34(1), 10–19. 10.1037/hea0000102

Cheval, B., Sarrazin, P., & Pelletier, L. (2014). Impulsive approach tendencies towards physical activity and sedentary behaviors, but not reflective intentions, prospectively predict non-exercise activity thermogenesis. Plos One, 9(12), e115238.

Cheval, B., Tipura, E., Burra, N., Frossard, J., Chanal, J., Orsholits, D., Radel, R., & Boisgontier, M. P. (2018). Avoiding sedentary behaviors requires more cortical resources than avoiding physical activity: An EEG study. Neuropsychologia, 119, 68–80. 10.1016/j.neuropsychologia.2018.07.029

Cheval, B., Zou, L., Maltagliati, S., Fessler, L., Owen, N., Falck, R. S., Yu, Q., Zhang, Z., & Dupuy, O. (2024). Intention–behaviour gap in physical activity: unravelling the critical role of the automatic tendency towards effort minimisation. British Journal of Sports Medicine. 10.1136/bjsports-2024-108144

Collins, D. L., Neelin, P., Peters, T. M., & Evans, A. C. (1994). Automatic 3D intersubject registration of MR volumetric data in standardized Talairach space. Journal of computer assisted tomography, 18(2), 192–205.

Conroy, D. E., & Berry, T. R. (2017). Automatic affective evaluations of physical activity. Exercise and Sport Sciences Reviews, 45(4), 230–237. 10.1249/JES.0000000000000120

Corbit, L. H., & Balleine, B. W. (2011). The general and outcome-specific forms of Pavlovian-instrumental transfer are differentially mediated by the nucleus accumbens core and shell. Journal of Neuroscience, 31(33), 11786–11794. 10.1523/JNEUROSCI.2711-11.2011

Craig, C. L., Marshall, A. L., Sjostrom, M., Bauman, A. E., Booth, M. L., Ainsworth, B. E., Pratt, M., Ekelund, U., Yngve, A., Sallis, J. F., & Oja, P. (2003). International physical activity questionnaire: 12-country reliability and validity. Medicine and Science in Sports and Exercise, 35(8), 1381–1395. 10.1249/01.MSS.0000078924.61453.FB

Crémers, J., Dessoullières, A., & Garraux, G. (2012). Hemispheric specialization during mental imagery of brisk walking. Human brain mapping, 33(4), 873–882. 10.1002/hbm.21255

Csajbók, Z., Sieber, S., Cullati, S., Cermakova, P., & Cheval, B. (2022). Physical activity partly mediates the association between cognitive function and depressive symptoms. Translational Psychiatry, 12(1), 414. 10.1038/s41398-022-02191-7

Daly, M., McMinn, D., & Allan, J. L. (2015). A bidirectional relationship between physical activity and executive function in older adults. Frontiers in human neuroscience, 8, 1044. 10.3389/fnhum.2014.01044

Davis, C., Katzman, D. K., Kaptein, S., Kirsh, C., Brewer, H., Kalmbach, K., Olmsted, M. F., Woodside, D. B., & Kaplan, A. S. (1997). The prevalence of high-level exercise in the eating disorders: etiological implications. Comprehensive psychiatry, 38(6), 321–326.

Ding, D., Lawson, K. D., Kolbe-Alexander, T. L., Finkelstein, E. A., Katzmarzyk, P. T., van Mechelen, W., Pratt, M., & Committee, L. P. A. S. E. (2016). The economic burden of physical inactivity: a global analysis of major non-communicable diseases. The Lancet, 388(10051), 1311–1324. 10.1016/S0140-6736(16)30383-X

Duckworth, A. L., & Kern, M. L. (2011). A meta-analysis of the convergent validity of self-control measures. Journal of Research in Personality, 45(3), 259–268. 10.1016/j.jrp.2011.02.004

Elliot, A. J., & Thrash, T. M. (2010). Approach and avoidance temperament as basic dimensions of personality. Journal of Personality, 78(3), 865–906. 10.1111/j.1467-6494.2010.00636.x

Epstein, R. A., Patai, E. Z., Julian, J. B., & Spiers, H. J. (2017). The cognitive map in humans: spatial navigation and beyond. Nature neuroscience, 20(11), 1504–1513. 10.1038/nn.4656

Farajzadeh, A., Goubran, M., Beehler, A., Cherkawi, N., Morrison, P., de Chanaleilles, M., Maltagliati, S., Cheval, B., Miller, M. W., Sheehy, L., & Boisgontier, M. (2023). Automatic approach-avoidance tendency toward physical activity, sedentary, and neutral stimuli as a function of age, explicit affective attitude, and intention to be active. Peer Community Journal, 3, e21. 10.24072/pcjournal.246

Faul, F., Erdfelder, E., Buchner, A., & Lang, A.-G. (2009). Statistical power analyses using G* Power 3.1: tests for correlation and regression analyses. Behavior Research Methods, 41(4), 1149–1160. 10.3758/BRM.41.4.1149

Frossard, J., & Renaud, O. (2019). The correlation structure of mixed effects models with crossed random effects in controlled experiments. *Preprint at* https://arxiv.org/abs/1903.10766.

Gottfried, J. A., O’Doherty, J., & Dolan, R. J. (2003). Encoding predictive reward value in human amygdala and orbitofrontal cortex. Science, 301(5636), 1104–1107. 10.1126/science.1087919

Griffiths, M., Szabo, A., & Terry, A. (2005). The exercise addiction inventory: a quick and easy screening tool for health practitioners. British Journal of Sports Medicine, 39(6), e30–e30.

Hannan, T. E., Moffitt, R. L., Neumann, D. L., & Kemps, E. (2019). Implicit approach– avoidance associations predict leisure-time exercise independently of explicit exercise motivation. *Sport*, Exercise, and Performance Psychology, 8(2), 210–222. 10.1037/spy0000145

Hernández, M. d. C. V., Cox, S. R., Kim, J., Royle, N. A., Maniega, S. M., Gow, A. J., Anblagan, D., Bastin, M. E., Park, J., & Starr, J. M. (2017). Hippocampal morphology and cognitive functions in community-dwelling older people: the Lothian Birth Cohort 1936. Neurobiology of aging, 52, 1–11. 10.1016/j.neurobiolaging.2016.12.012

Jackson, T., Gao, X., & Chen, H. (2014). Differences in neural activation to depictions of physical exercise and sedentary activity: an fMRI study of overweight and lean Chinese women. International Journal of Obesity, 38(9), 1180. 10.1038/ijo.2013.245

Jacobsen, L. K., Gore, J. C., Skudlarski, P., Lacadie, C. M., Jatlow, P., & Krystal, J. H. (2002). Impact of intravenous nicotine on BOLD signal response to photic stimulation. Magnetic resonance imaging, 20(2), 141–145.

Jenkinson, M., Beckmann, C. F., Behrens, T. E., Woolrich, M. W., & Smith, S. M. (2012). Fsl. Neuroimage, 62(2), 782–790. 10.1016/j.neuroimage.2011.09.015

Klur, S., Muller, C., Pereira de Vasconcelos, A., Ballard, T., Lopez, T., Galani, R., Certa, U., & Cassel, J. C.. (2009). Hippocampal-dependent spatial memory functions might be lateralized in rats: An approach combining gene expression profiling and reversible inactivation. Hippocampus, 19(9), 800–816. 10.1002/hipo.20562

Knutson, B., Adams, C. M., Fong, G. W., & Hommer, D. (2001). Anticipation of increasing monetary reward selectively recruits nucleus accumbens. Journal of Neuroscience, 21(16), RC159. 10.1523/JNEUROSCI.21-16-j0002.2001

Krieglmeyer, R., & Deutsch, R. (2010). Comparing measures of approach–avoidance behaviour: The manikin task vs. two versions of the joystick task. Cognition and Emotion, 24(5), 810–828. 10.1080/02699930903047298

Kullmann, S., Giel, K. E., Hu, X., Bischoff, S. C., Teufel, M., Thiel, A., Zipfel, S., & Preissl, H. (2014). Impaired inhibitory control in anorexia nervosa elicited by physical activity stimuli. Social cognitive and affective neuroscience, 9(7), 917–923. 10.1093/scan/nst070

Kuznetsova, A., Brockhoff, P. B., & Christensen, R. H. B. (2015). lmerTest Package: tests in linear mixed effects models. Journal of statistical software, 82(13). 10.18637/jss.v082.i13

Luciani, J. (2015). Why 80 Percent of New Year’s Resolutions Fail. Retrieved from https://health.usnews.com/health-news/blogs/eat-run/articles/2015-12-29/why-80-percent-of-new-years-resolutions-fail.

Maltagliati, S., Fessler, L., Yu, Q., Zhang, Z., Chen, Y., Dupuy, O., Falck, R., Owen, N., Zou, L., & Cheval, B. (2024). Effort Minimization: A Permanent, Dynamic, and Surmountable Influence on Physical Activity Journal of Sport and Health Science, 100971. 10.1016/j.jshs.2024.100971

Maltagliati, S., Raichlen, D. A., Rhodes, R. E., & Cheval, B. (2024). Closing the intention-behavior gap in physical activity: the moderating effect of individual differences in the valuation of physical effort. SportArxiv. 10.51224/SRXIV.375

Moffitt, R. L., Kemps, E., Hannan, T. E., Neumann, D. L., Stopar, S. P., & Anderson, C. J. (2019). Implicit approach biases for physically active lifestyle cues. International Journal of Sport and Exercise Psychology, 18(6), 833–849. 10.1080/1612197X.2019.1581829

Nash, J. C., & Varadhan, R. (2011). Unifying optimization algorithms to aid software system users: optimx for R. Journal of statistical software, 43(9). 10.18637/jss.v043.i09

Nelder, J. A., & Mead, R. (1965). A simplex method for function minimization. The Computer Journal, 7(4), 308–313. 10.1093/comjnl/7.4.308

Oldfield, R. C. (1971). The assessment and analysis of handedness: the Edinburgh inventory. Neuropsychologia, 9(1), 97–113. 10.1016/0028-3932(71)90067-4

Parma, J., Bacelar, M., Cabral, D., Recker, R., Renaud, O., Sander, D., Krigolson, O., Miller, M., Cheval, B., & Boisgontier, M. (2023). Relationship between reward-related brain activity and opportunities to sit. Cortex, 167, 197–217. 10.1016/j.cortex.2023.06.011

Patenaude, B., Smith, S. M., Kennedy, D. N., & Jenkinson, M. (2011). A Bayesian model of shape and appearance for subcortical brain segmentation. Neuroimage, 56(3), 907–922. 10.1016/j.neuroimage.2011.02.046

Phaf, R. H., Mohr, S. E., Rotteveel, M., & Wicherts, J. M. (2014). Approach, avoidance, and affect: a meta-analysis of approach-avoidance tendencies in manual reaction time tasks. Frontiers in Psychology, 5, 378. 10.3389/fpsyg.2014.00378

Pool, E., Brosch, T., Delplanque, S., & Sander, D. (2016). Attentional bias for positive emotional stimuli: a meta-analytic investigation. Psychological Bulletin, 142, 79–106. 10.1037/bul0000026

Powell, M. J. (2009). The BOBYQA algorithm for bound constrained optimization without derivatives. Cambridge NA Report NA2009/06, University of Cambridge, Cambridge, 26–46.

Prévost, C., Liljeholm, M., Tyszka, J. M., & O’Doherty, J. P. (2012). Neural correlates of specific and general Pavlovian-to-Instrumental Transfer within human amygdalar subregions: a high-resolution fMRI study. Journal of Neuroscience, 32(24), 8383–8390. 10.1523/JNEUROSCI.6237-11.2012

Prévost, C., Pessiglione, M., Météreau, E., Cléry-Melin, M.-L., & Dreher, J.-C. (2010). Separate valuation subsystems for delay and effort decision costs. Journal of Neuroscience, 30(42), 14080–14090. 10.1523/JNEUROSCI.2752-10.2010

Roesch, M. R., & Olson, C. R. (2004). Neuronal activity related to reward value and motivation in primate frontal cortex. Science, 304(5668), 307–310. 10.1126/science.1093223

Rolls, E. T., Huang, C.-C., Lin, C.-P., Feng, J., & Joliot, M. (2020). Automated anatomical labelling atlas 3. Neuroimage, 206, 116189. 10.1016/j.neuroimage.2019.116189

Rougier, M., Muller, D., Ric, F., Alexopoulos, T., Batailler, C., Smeding, A., & Aubé, B. (2018). A new look at sensorimotor aspects in approach/avoidance tendencies: The role of visual whole-body movement information. Journal of Experimental Social Psychology, 76, 42–53. 10.1016/j.jesp.2017.12.004

Sabia, S., Dugravot, A., Dartigues, J.-F., Abell, J., Elbaz, A., Kivimäki, M., & Singh-Manoux, A. (2017). Physical activity, cognitive decline, and risk of dementia: 28 year follow-up of Whitehall II cohort study. British Medical Journal, 357, j2709. 10.1136/bmj.j2709

Schultz, W., Tremblay, L., & Hollerman, J. R. (2000). Reward processing in primate orbitofrontal cortex and basal ganglia. Cerebral cortex, 10(3), 272–283.

Smith, S. M., Jenkinson, M., Woolrich, M. W., Beckmann, C. F., Behrens, T. E., Johansen-Berg, H., Bannister, P. R., De Luca, M., Drobnjak, I., & Flitney, D. E. (2004). Advances in functional and structural MR image analysis and implementation as FSL. Neuroimage, 23, S208–S219. 10.1016/j.neuroimage.2004.07.051

Smith, S. M., & Nichols, T. E. (2009). Threshold-free cluster enhancement: addressing problems of smoothing, threshold dependence and localisation in cluster inference. Neuroimage, 44(1), 83–98. 10.1016/j.neuroimage.2008.03.061

Strack, F., & Deutsch, R. (2004). Reflective and impulsive determinants of social behavior. Personality and Social Psychology Review, 8(3), 220–247. 10.1207/s15327957pspr0803_1

Strain, T., Flaxman, S., Guthold, R., Semenova, E., Cowan, M., Riley, L. M., Bull, F. C., & Stevens, G. A. (2024). National, regional, and global trends in insufficient physical activity among adults from 2000 to 2022: a pooled analysis of 507 population-based surveys with 5· 7 million participants. The Lancet Global Health, 12(8), E1232–E1243. 10.1016/S2214-109X(24)00150-5

Team, R. C. (2019). R Core Team. R: A language and environment for statistical computing. In: Vienna, Austria. https://www.R-project.org/.

WHO. (2019). Global action plan on physical activity 2018-2030: more active people for a healthier world. Retrieved from https://apps.who.int/iris/bitstream/handle/10665/272722/9789241514187-eng.pdf.

WHO. (2020). WHO guidelines on physical activity and sedentary behaviour. Retrieved from https://www.who.int/publications/i/item/9789240015128.

Woolrich, M. W., Jbabdi, S., Patenaude, B., Chappell, M., Makni, S., Behrens, T., Beckmann, C., Jenkinson, M., & Smith, S. M. (2009). Bayesian analysis of neuroimaging data in FSL. Neuroimage, 45(1), S173–S186. 10.1016/j.neuroimage.2008.10.055

Zandbelt, B. B., & Vink, M. (2010). On the role of the striatum in response inhibition. PloS one, 5(11), e13848.

