## Supplemental Tables for "Neural correlates of approach and avoidance tendencies toward physical activity and sedentary stimuli: An fMRI study"

**Supplemental materials**

**S1. Measures of the dispositional and situational moderators**

**S2. Results of the exploratory analyses for reaction times**

**S3. Results of the exploratory analyses for errors**

**S4. Neural Activity Associated with the Avoidance of Sedentary Stimuli: Avoid Sedentary > Avoid Neutral (HN4).**

**S5. Detailed coordinates of the clusters presented in this section.**

**S1. Measures of the dispositional and situational moderators**

**Usual level of physical activity.** Commuting and leisure time physical activity was assessed using an adapted version of the International Physical Activity Questionnaire. Specifically, the first set of questions focused on the sports-related physical activity, the second on commuter-related physical activity (i.e., walking and cycling), and the last set of questions focused on leisure-time related physical activity (i.e., walking, moderate physical activity, vigorous physical activity). Here we created a score that is the sum of all the time spent in these different physical activities.

**Craving for sedentary behaviors.** Sedentary craving was assessed with the following item: “At this moment, to what extent do you feel the desire (urge) to engage in sedentary activities (e.g., sitting, resting, watching TV)?”. Participants respond on a scale from 1 “no desire at all” to 7 “very strong desire”.

**Craving for physical activity.** Physical activity craving was assessed with the following item: “At this moment, to what extent do you feel the desire (urge) to engage in physical activity (e.g., running, exercising, moving)?”. Participants respond on a scale from 1 “no desire at all” to 7 “very strong desire”.

**S2. Results of the exploratory analyses for reaction times**

**Usual level of physical activity**

*Results of the linear mixed-effects models predicting the reaction times as a function of the type of action (approach vs. avoidance) and of the type of stimuli (physical activity vs. neutral vs. physical inactivity) and usual level of physical activity*

| **Moderation by the usual level of physical activity**  **(N = 40)** | b (CI) | *p* | |
| --- | --- | --- | --- |
| **Fixed Effects** |  |  | |
| Intercept | **665.7 (627.2;704.2)** | | **<.001** |
| **Stimuli (ref. physical activity)** |  | |  |
| Neutral | 2.7 (-11.8;17.3) | | .712 |
| Sedentary | 7.9 (-6.0;18.93) | | .267 |
| **Action (ref. approach)** |  | |  |
| Avoidance | -6.9 (-23.6;21.9) | | .415 |
| **Stimuli (ref. physical activity) x Action (ref. approach)** |  | |  |
| Avoidance x neutral | -4.1 (-22.5;14.2) | | .661 |
| Avoidance x Sedentary | -1.7 (-20.1;16.6) | | .852 |
| **Usual level of physical activity** |  |  | |
| Usual level of physical activity | 13.3 (-21.2;47.8) | .454 | |
| Usual level of physical activity x  Sedentary stimuli | -6.0 (-19.2;7.2) | .373 | |
| Usual level of physical activity x  Neutral stimuli | -2.4 (-15.4;10.6) | .718 | |
| Usual level of physical activity x  Avoidance | -7.6 (-24.2;8.9) | .369 | |
| Usual level of physical activity x  neutral stimuli x Avoidance | -1.4 (-19.8;16.9) | .877 | |
| Usual level of physical activity x  Sedentary x Avoidance | 9.1 (-9.2;27.5) | .330 | |
| **Covariates** |  |  | |
| Age | -12.7 (-48.27;21.8) | .476 | |
| Sex | -3.9 (-73.27;70.3) | .918 | |
| BMI | -0.7 (-31.21;35.0) | .968 | |
| **Random Effects** |  |  | |
| **Participants** |  |  | |
| Intercept | 10877.52 | | |
| Stimuli sedentary | 55.82 | | |
| Stimuli neutral | 10.32 | | |
| Action Avoid | 1130.10 | | |
| Corr. (Intercept, stimuli sedentary) | 0.09 | | |
| Corr. (Intercept, stimuli neutral) | 0.96 | | |
| Corr. (Intercept, action avoidance) | -0.29 | | |
| Corr. (Stimuli sedentary; stimuli neutral) | -0.19 | | |
| Corr. (Stimuli sedentary; action avoidance) | 0.93 | | |
| Corr. (Stimuli neutral; action avoidance) | -0.55 | | |
| **Stimuli** |  |  | |
| Intercept | 95.08 | | |
| Residual | 28558.12 | | |
| R^2^ | Conditional .006  Marginal =.280 | | |

*Notes*. 95CI = confidence intervals at 95%.

**Craving for sedentary behaviors**

*Results of the linear mixed-effects models predicting the reaction times as a function of the type of action (approach vs. avoidance) and of the type of stimuli (physical activity vs. neutral vs. sedentary) and craving for sedentary behaviors*

| **Moderation by the craving for sedentary behaviors**  **(N = 40)** | b (CI) | *p* | |
| --- | --- | --- | --- |
| **Fixed Effects** |  |  | |
| Intercept | **664.6 (626.1;703.2)** | | **<.001** |
| **Stimuli (ref. physical activity)** |  | |  |
| Neutral | 2.8 (-11.7;17.3) | | .706 |
| Sedentary | 8.1 (-5.7;21.9) | | .251 |
| **Action (ref. approach)** |  | |  |
| Avoidance | -7.0 (-23.5;9.5) | | .407 |
| **Stimuli (ref. physical activity) x Action (ref. approach)** |  | |  |
| Avoidance x Neutral | -4.2 (-22.5;14.2) | | .565 |
| Avoidance x Sedentary | -2.1 (-20.4;16.3) | | .826 |
| **Craving for sedentary behaviors** |  |  | |
| Craving for sedentary behaviors | -3.2 (-37.0;30.5) | .852 | |
| Craving for sedentary behaviors x  Sedentary stimuli | **-22.0 (-35.0;-9.0)** | **<.001** | |
| Craving for sedentary behaviors x  Neutral stimuli | -5.1 (-18.1;8.0) | .449 | |
| Craving for sedentary behaviors x  Avoidance | -14.45 (-30.9;2.0) | .089 | |
| Craving for sedentary behaviors x  neutral stimuli x Avoidance | 3.0 (-153;21.36) | .747 | |
| Craving for sedentary behaviors x  Sedentary stimuli x Avoidance | 15.3 (-3.0;33.65) | .102 | |
| **Covariates** |  |  | |
| Age | -11.4 (-45.60;20.67) | .514 | |
| Sex | -0.14 (-55.55;85.92) | .997 | |
| BMI | 3.0 (-27.83;39.00) | .864 | |
| **Random Effects** |  |  | |
| **Participants** |  |  | |
| Intercept | 10979.23 | | |
| Stimuli sedentary | 11.29 | | |
| Stimuli neutral | 9.07 | | |
| Action Avoid | 1092.45 | | |
| Corr. (Intercept, stimuli sedentary) | -0.30 | | |
| Corr. (Intercept, stimuli neutral) | 0.84 | | |
| Corr. (Intercept, action avoidance) | -0.33 | | |
| Corr. (Stimuli sedentary; stimuli neutral) | -0.77 | | |
| Corr. (Stimuli sedentary; action avoidance) | 1.00 | | |
| Corr. (Stimuli neutral; action avoidance) | -0.79 | | |
| **Stimuli** |  |  | |
| Intercept | 95.40 | | |
| Residual | 28524.59 | | |
| R^2^ | Conditional .012  Marginal =.280 | | |

*Notes*. 95CI = confidence intervals at 95%.

**Figure S1.**

*Results of the linear mixed-effects models. A. Region of significance of the difference in reaction times to respond to sedentary relative to physical activity stimuli as function of the level of craving for sedentary behaviors.* *B. Prediction of the response times as a function of the type of stimuli and the level of craving for sedentary behaviors.*

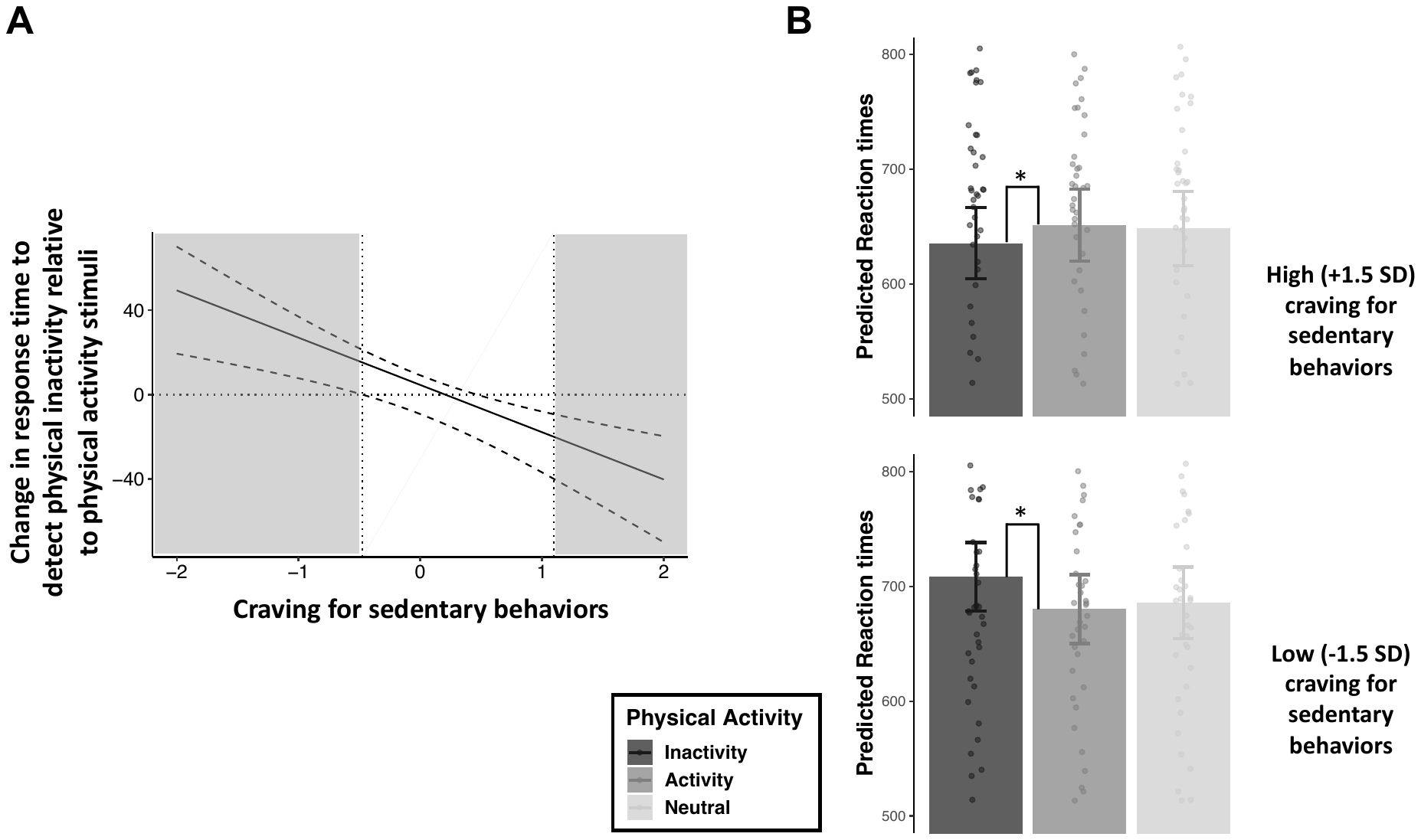

Notes. A. A positive effect (i.e., grey area on the left) indicates that participants were significantly slower to respond to sedentary relative to physical activity stimuli, while a negative effect (i.e., grey area on the right) indicates the contrary – participants were significantly quicker to respond to sedentary relative to physical activity stimuli. Craving for sedentary behaviors are standardized. B. Dots representing mean response times for each participant as a function of the type of stimuli (i.e., physical activity vs. sedentary vs. neutral stimuli). Errors bars represent the standard errors around the mean.

**Craving for physical activity**

*Results of the linear mixed-effects models predicting the reaction times as a function of the type of action (approach vs. avoidance) and of the type of stimuli (physical activity vs. neutral vs. sedentary) and craving for physical activity*

| **Moderation by the craving for physical activity**  **(N = 40)** | b (CI) | *p* | |
| --- | --- | --- | --- |
| **Fixed Effects** |  |  | |
| Intercept | **665.8 (627.0;704.5)** | | **<.001** |
| **Stimuli (ref. physical activity)** |  | |  |
| Neutral | 2.8 (-11.7;17.3) | | .707 |
| Sedentary | 8.0 (-6.0;21.9) | | .264 |
| **Action (ref. approach)** |  | |  |
| Avoidance | -6.9 (-23.6;9.8) | | .418 |
| **Stimuli (ref. physical activity) x Action (ref. approach)** |  | |  |
| Avoidance x neutral | -4.2 (-22.5;14.2) | | .656 |
| Avoidance x Sedentary | -1.9 (-20.2;16.5) | | .842 |
| **Craving for physical activity** |  |  | |
| Craving for physical activity | -8.8 (-43.2;25.7) | .919 | |
| Craving for physical activity x  Sedentary stimuli | 9.5 (-3.7;22.7) | .159 | |
| Craving for physical activity x  Neutral stimuli | 8.2 (-4.9;21.2) | .218 | |
| Craving for physical activity x  Avoidance | 2.5 (-14.2;19.2) | .773 | |
| Craving for physical activity x  neutral stimuli x Avoidance | -8.0 (-26.3;10.3) | .392 | |
| Craving for physical activity x  Sedentary stimuli x Avoidance | -1.2 (-19.6;17.2) | .899 | |
| **Covariates** |  |  | |
| Age | -13.3 (-48.3;20.67) | .460 | |
| Sex | -4.1 (-78.6;85.92) | .914 | |
| BMI | 1.8 (-33.1;39.00) | .918 | |
| **Random Effects** |  |  | |
| **Participants** |  |  | |
| Intercept | 10976.35 | | |
| Stimuli sedentary | 56.98 | | |
| Stimuli neutral | 8.9 | | |
| Action Avoid | 1158.77 | | |
| Corr. (Intercept, stimuli sedentary) | 0.09 | | |
| Corr. (Intercept, stimuli neutral) | 0.98 | | |
| Corr. (Intercept, action avoidance) | -0.30 | | |
| Corr. (Stimuli sedentary; stimuli neutral) | -0.13 | | |
| Corr. (Stimuli sedentary; action avoidance) | 0.93 | | |
| Corr. (Stimuli neutral; action avoidance) | -0.50 | | |
| **Stimuli** |  |  | |
| Intercept | 94.44 | | |
| Residual | 28549.61 | | |
| R^2^ | Conditional .005  Marginal =.281 | | |

*Notes*. 95CI = confidence intervals at 95%.

**S3. Results of the exploratory analyses for errors**

**Usual level of physical activity**

*Results of the logistic mixed-effects models predicting the risk of error in the approach-avoidance task as a function of action type (approach vs. avoidance) and stimuli type (physical activity vs. neutral vs. sedentary), and usual physical activity.*

| **N = 40** | OR (CI) | *p* |
| --- | --- | --- |
| **Fixed Effects** |  |  |
| Intercept | **0.06 (0.04;0.08)** | **<.001** |
| **Stimuli (ref. physical activity)** |  |  |
| Neutral | 0.80 (0.59;1.09) | .151 |
| Sedentary | 0.85 (0.63;1.15) | .294 |
| **Action (ref. approach)** |  |  |
| Avoidance | 0.73 (0.52;1.03) | .070 |
| **Stimuli (ref. physical activity) x Action (ref. approach)** |  |  |
| Avoidance x neutral | **1.57 (1.01;2.44)** | **.045** |
| Avoidance x Sedentary | **1.64 (1.06;2.54)** | **.025** |
| **Usual level of physical activity** |  |  |
| Usual level of physical activity | 1.03 (0.72;1.45) | .887 |
| Usual level of physical activity x  Sedentary stimuli | 1.02 (0.74;1.40) | .906 |
| Usual level of physical activity x  Neutral stimuli | 0.96 (0.70;1.33) | .814 |
| Usual level of physical activity x  Avoidance | 0.91 (0.64;1.28) | .575 |
| Usual level of physical activity x  neutral stimuli x Avoidance | 1.05 (0.66;1.66) | .850 |
| Usual level of physical activity x  Sedentary x Avoidance | 1.10 (0.70;1.74) | .669 |
| **Covariates** |  |  |
| Age | 1.04 (0.77;1.40) | .808 |
| Sex | 0.95 (0.49;1.83) | . 881 |
| BMI | 0.91 (0.66;1.24) | .542 |
| **Random Effects** |  |  |
| **Participants** |  |  |
| Intercept | 0.68 | |
| Stimuli sedentary | 0.01 | |
| Stimuli neutral | 0.02 | |
| Action Avoid | 0.10 | |
| Corr. (Intercept, stimuli sedentary) | 0.79 | |
| Corr. (Intercept, stimuli neutral) | 0.99 | |
| Corr. (Intercept, action avoidance) | -0.39 | |
| Corr. (Stimuli sedentary; stimuli neutral) | 0.71 | |
| Corr. (Stimuli sedentary; action avoidance) | -0.87 | |
| Corr. (Stimuli neutral; action avoidance) | -0.27 | |
| **Stimuli** |  | |
| Intercept | Null | |
| R^2^ | Conditional .006  Marginal =.192 | |

*Notes*. OR= odds ratio; 95CI = confidence intervals at 95%. Note that the models estimated a null variance for the random intercept of the stimuli. The models with or without this parameter lead to consistent results. *Two participants were not included in the analyses because they were an issue regarding the recording of their behavioral data.

**Craving for sedentary behaviors**

*Results of the logistic mixed-effects models predicting the risk of error in the approach-avoidance task as a function of action type (approach vs. avoidance) and stimuli type (physical activity vs. neutral vs. sedentary), and craving for sedentary behaviors.*

| **N = 40** | OR (CI) | *p* |
| --- | --- | --- |
| **Fixed Effects** |  |  |
| Intercept | **0.06 (0.04;0.08)** | **<.001** |
| **Stimuli (ref. physical activity)** |  |  |
| Neutral | 0.82 (0.60;1.11) | .193 |
| Sedentary | 0.82 (0.61;1.11) | .224 |
| **Action (ref. approach)** |  |  |
| Avoidance | 0.74 (0.53;1.03) | .069 |
| **Stimuli (ref. physical activity) x Action (ref. approach)** |  |  |
| Avoidance x neutral | **1.49 (0.96;2.31)** | **.018** |
| Avoidance x Sedentary | **1.70 (1.10;2.63)** | **.079** |
| **Craving for sedentary behaviors** |  |  |
| Craving for sedentary behaviors | 0.80 (0.57;1.13) | .180 |
| Craving for sedentary behaviors x  Sedentary stimuli | 0.84 (0.61;1.15) | .245 |
| Craving for sedentary behaviors x  Neutral stimuli | 1.03 (0.75;1.42) | .854 |
| Craving for sedentary behaviors x  Avoidance | 1.16 (0.82;1.65) | .345 |
| Craving for sedentary behaviors x  neutral stimuli x Avoidance | 0.73 (0.46;1.15) | .125 |
| Craving for sedentary behaviors x  Sedentary stimuli x Avoidance | 1.26 (0.80;1.98) | .276 |
| **Covariates** |  |  |
| Age | 1.04 (0.77;1.40) | .800 |
| Sex | 0.94 (0.49;1.82) | .864 |
| BMI | 0.92 (0.67;1.26) | .593 |
| **Random Effects** |  |  |
| **Participants** |  |  |
| Intercept | 0.64 | |
| Stimuli sedentary | 0.01 | |
| Stimuli neutral | 0.02 | |
| Action Avoid | 0.08 | |
| Corr. (Intercept, stimuli sedentary) | 0.59 | |
| Corr. (Intercept, stimuli neutral) | 0.98 | |
| Corr. (Intercept, action avoidance) | -0.31 | |
| Corr. (Stimuli sedentary; stimuli neutral) | 0.42 | |
| Corr. (Stimuli sedentary; action avoidance) | -0.95 | |
| Corr. (Stimuli neutral; action avoidance) | -0.11 | |
| **Stimuli** |  | |
| Intercept | Null | |
| R^2^ | Conditional .021  Marginal =.196 | |

*Notes*. OR= odds ratio; 95CI = confidence intervals at 95%. Note that the models estimated a null variance for the random intercept of the stimuli. The models with or without this parameter lead to consistent results. *Two participants were not included in the analyses because they were an issue regarding the recording of their behavioral data.

**Craving for physical activity**

*Results of the logistic mixed-effects models predicting the risk of error in the approach-avoidance task as a function of action type (approach vs. avoidance) and stimuli type (physical activity vs. neutral vs. sedentary), and craving for physical activity.*

| **N = 40** | OR (CI) | *p* |
| --- | --- | --- |
| **Fixed Effects** |  |  |
| Intercept | **0.6 (0.04;0.08)** | **<.001** |
| **Stimuli (ref. physical activity)** |  |  |
| Neutral | 0.78 (0.57;1.06) | .117 |
| Sedentary | 0.85 (0.62;1.15) | .289 |
| **Action (ref. approach)** |  |  |
| Avoidance | 0.71 (0.51;1.00) | .052 |
| **Stimuli (ref. physical activity) x Action (ref. approach)** |  |  |
| Avoidance x neutral | **1.67 (1.07;2.61)** | **.024** |
| Avoidance x Sedentary | **1.65 (1.06;2.59)** | **.027** |
| **Craving for physical activity** |  |  |
| Craving for physical activity | 0.96 (0.69;1.34) | .799 |
| Craving for physical activity x  Sedentary stimuli | 0.83 (0.62;1.12) | .218 |
| Craving for physical activity x  Neutral stimuli | 0.86 (0.64;1.16) | .331 |
| Craving for physical activity x  Avoidance | 0.77 (0.55;1.07) | .119 |
| Craving for physical activity x  neutral stimuli x Avoidance | 1.43 (0.94;2.19) | .097 |
| Craving for physical activity x  Sedentary stimuli x Avoidance | 1.11 (0.73;1.70) | .620 |
| **Covariates** |  |  |
| Age | 1.04 (0.74;1.33) | .948 |
| Sex | 0.94 (0.53;1.89) | .997 |
| BMI | 0.92 (0.69;1.25) | .610 |
| **Random Effects** |  |  |
| **Participants** |  |  |
| Intercept | 0.67 | |
| Stimuli sedentary | 0.01 | |
| Stimuli neutral | 0.02 | |
| Action Avoid | 0.10 | |
| Corr. (Intercept, stimuli sedentary) | 0.40 | |
| Corr. (Intercept, stimuli neutral) | 1.00 | |
| Corr. (Intercept, action avoidance) | -0.44 | |
| Corr. (Stimuli sedentary; stimuli neutral) | 0.40 | |
| Corr. (Stimuli sedentary; action avoidance) | -1.00 | |
| Corr. (Stimuli neutral; action avoidance) | -0.43 | |
| **Stimuli** |  | |
| Intercept | Null | |
| R^2^ | Conditional .020  Marginal =.195 | |

*Notes*. OR= odds ratio; 95CI = confidence intervals at 95%. Note that the models estimated a null variance for the random intercept of the stimuli. The models with or without this parameter lead to consistent results. *Two participants were not included in the analyses because they were an issue regarding the recording of their behavioral data.

**S4. Neural Activity Associated with the Avoidance of Sedentary Stimuli: *Avoid Sedentary > Avoid Neutral (HN4).***

Brain activations when avoiding sedentary vs. neutral stimuli, corrected for multiple comparisons (whole-brain voxel-wise *p*<.05 FDR, k>10 voxels). The color bar represents the statistical T value. SPL: superior parietal lobule; TOC: temporo-occipital cortex; V1: primary visual cortex; V3/4: associative visual cortex. L: left hemisphere; R: right hemisphere.

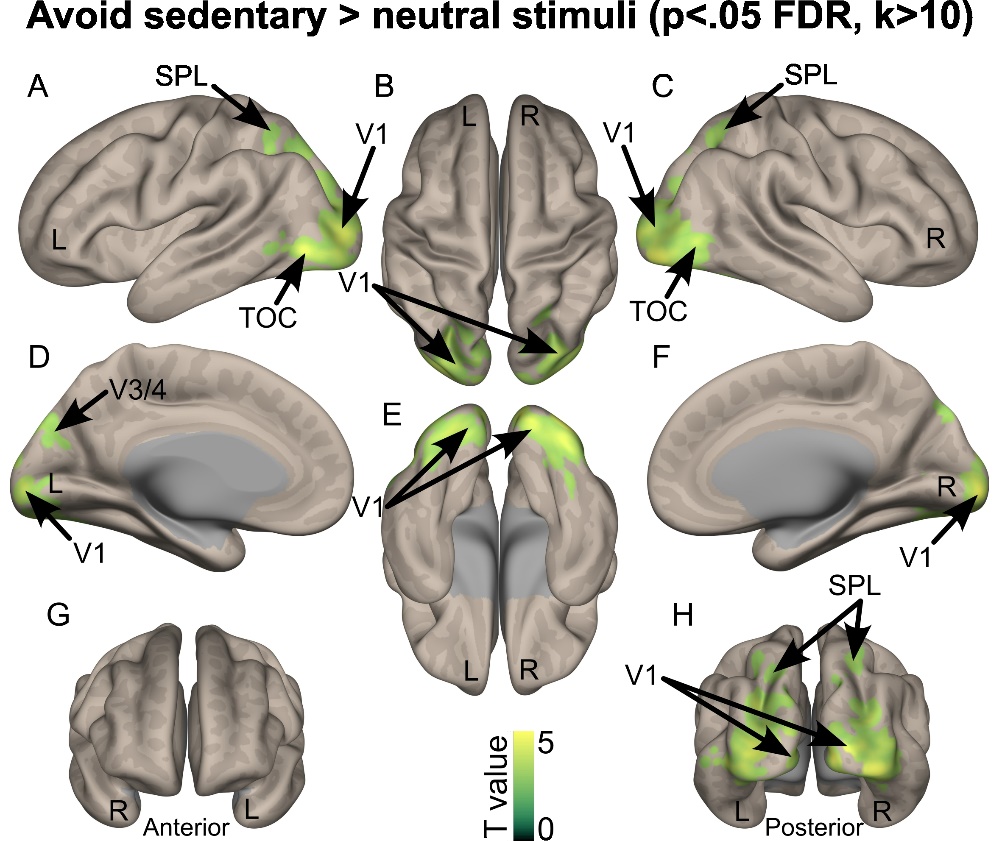

**S5. Detailed coordinates of the clusters presented in this section.**

**MNI coordinates for the neuroimaging results of the Approach vs Avoid sedentary stimuli contrast***.*

| **Region** | **Hemisphere** | **Cluster size** | **T value** | **MNI X** | **MNI Y** | **MNI Z** |
| --- | --- | --- | --- | --- | --- | --- |
| Fusiform cortex | R | 1594 | 5,65 | 28 | -56 | -8 |
| Mid Occipital cortex (V1) | L |  | 5,34 | -14 | -94 | 0 |
| Fusiform cortex | R |  | 5,24 | 28 | -76 | -12 |
| Mid Occipital cortex (V1) | R |  | 4,73 | 18 | -84 | -10 |
| Inf Occipital cortex (V2/3) | R |  | 4,69 | 36 | -80 | -12 |
| Parahippocampal gyrus | L | 159 | 4,58 | -26 | -44 | -8 |
| Fusiform cortex | L | 31 | 4,34 | -36 | -40 | -22 |
| Mid Occipital cortex | L | 92 | 4,20 | -30 | -78 | 24 |
| Mid Occipital cortex | L |  | 3,71 | -26 | -84 | 14 |
| Mid Occipital cortex | L |  | 3,48 | -38 | -80 | 18 |
| Mid Occipital cortex | R | 60 | 4,15 | 32 | -76 | 20 |
| Thalamus | R | 12 | 3,76 | 24 | -26 | -6 |
| Parahippocampal gyrus | L | 11 | 3,64 | -20 | -32 | -8 |

V1: primary visual cortex; V2/3: secondary visual cortex. MNI: Montreal Neurological Institute.

**MNI coordinates for the neuroimaging results of the Avoid vs Approach sedentary stimuli contrast***.*

| **Region** | **Hemisphere** | **Cluster size** | **T value** | **MNI X** | **MNI Y** | **MNI Z** |
| --- | --- | --- | --- | --- | --- | --- |
| SMA | R | 1356 | 4,85 | 2 | -18 | 66 |
| Mid Cingulate cortex | R |  | 4,73 | 8 | -32 | 54 |
| SMA | R |  | 4,72 | 10 | -24 | 58 |
| Mid Cingulate cortex | R |  | 4,71 | 6 | -16 | 46 |
| Primary motor cortex | R |  | 4,61 | 28 | -30 | 60 |
| Insula | R | 213 | 4,49 | 46 | 8 | 4 |
| IFG tri | R |  | 3,94 | 46 | 24 | 2 |
| IFG oper | R |  | 3,92 | 46 | 16 | 0 |
| IFG orb | R |  | 3,66 | 54 | 22 | -4 |
| DLPFC | R | 227 | 4,47 | 28 | 44 | 18 |
| DLPFC | R |  | 4,25 | 34 | 50 | 18 |
| DLPFC | R |  | 3,71 | 26 | 38 | 26 |
| Primary motor cortex | L | 275 | 4,35 | -20 | -26 | 66 |
| S1 | L |  | 4,32 | -22 | -34 | 56 |
| S1 | L |  | 3,57 | -32 | -42 | 58 |
| STG post | R | 94 | 4,19 | 48 | -24 | 14 |
| STS mid | R | 209 | 4,16 | 50 | -32 | 0 |
| STG post | R |  | 4,13 | 54 | -24 | -2 |
| Temporal pole | R | 41 | 4,11 | 46 | 22 | -18 |
| Primary motor cortex | L | 182 | 4,07 | -50 | -8 | 46 |
| Pre primary motor cortex | R | 122 | 4,06 | 44 | -6 | 40 |
| IFG oper | L | 59 | 3,95 | -50 | 6 | 2 |
| Insula | L | 46 | 3,95 | -42 | 4 | 10 |
| Frontal gyrus mid | L | 61 | 3,91 | -28 | 52 | 6 |
| Cingulate cortex mid | R | 24 | 3,91 | 10 | 18 | 42 |
| Cingulate cortex mid | R | 42 | 3,90 | 10 | -4 | 38 |
| Heschl gyrus | L | 70 | 3,85 | -34 | -26 | 6 |
| Planum temporale | L |  | 3,84 | -42 | -22 | 8 |
| Cerebellum lobule 6 | L | 14 | 3,70 | -26 | -44 | -34 |
| Thalamus | L | 26 | 3,68 | -22 | -12 | 14 |
| Putamen | L |  | 3,31 | -28 | -8 | 10 |
| Insula | L | 19 | 3,65 | 44 | -6 | 10 |

SMA: supplementary motor area; IFG: inferior frontal gyrus (tri: *pars triangularis*, oper: *pars opercularis*, orb: *pars orbitalis*); DLPFC: dorsolateral prefrontal cortex; S1: primary somatosensory cortex; STG: superior temporal gyrus; STS: superior temporal sulcus; post: posterior.

**Table 6. MNI coordinates for the neuroimaging results of the Avoid sedentary stimuli vs Avoid physical activity contrast***.*

| **Region** | **Hemisphere** | **Cluster size** | **T value** | **MNI X** | **MNI Y** | **MNI Z** |
| --- | --- | --- | --- | --- | --- | --- |
| STG mid | R | 614 | 5,01 | 50 | -6 | -6 |
| STG mid | R |  | 4,84 | 52 | -18 | -6 |
| STS post | R |  | 4,50 | 50 | -34 | -2 |
| STG post | R |  | 4,47 | 66 | -44 | 16 |
| Thalamus | R | 145 | 4,80 | 4 | -12 | 4 |
| Thalamus | R |  | 4,01 | 12 | -26 | -2 |
| Substania nigra | R |  | 3,60 | 14 | -18 | -8 |
| SMA | L | 78 | 4,72 | -14 | -24 | 66 |
| Cerebellum lobule 4-5 | R | 80 | 4,37 | 10 | -56 | -20 |
| Cerebellum lobule 4-6 | R |  | 3,82 | 10 | -44 | -12 |
| Globus pallidus | L | 48 | 4,34 | -20 | 0 | -2 |
| Putamen | L |  | 3,57 | -32 | 0 | -4 |
| Temporal pole | L | 54 | 4,26 | -50 | 12 | -20 |
| Globus pallidus | R | 118 | 4,20 | 30 | -10 | -4 |
| Globus pallidus | R |  | 4,05 | 20 | -6 | 2 |
| Putamen | R |  | 3,81 | 28 | -8 | 6 |
| Cerebellum Crus 1 | L | 58 | 4,10 | -18 | -68 | -28 |
| Cerebellum lobule 6 | R | 12 | 3,96 | 16 | -70 | -26 |
| STG post | L | 21 | 3,93 | -54 | -46 | 20 |
| DLPFC | R | 51 | 3,91 | 28 | 48 | 20 |
| Insula | R | 30 | 3,87 | -34 | 8 | -8 |
| Cingulate cortex mid | L | 13 | 3,86 | 8 | -18 | 44 |
| MTG ant | L | 14 | 3,82 | -56 | -6 | -24 |
| Angular gyrus | L | 16 | 3,61 | -50 | -66 | 24 |

STG: superior temporal gyrus; STS: superior temporal sulcus (post: posterior); SMA: supplementary motor area; DLPFC: dorsolateral prefrontal cortex; MTG: middle temporal gyrus (ant: anterior).

**Table 7. MNI coordinates for the neuroimaging results of the Avoid sedentary stimuli vs Avoid Neutral stimuli contrast***.*

| **Region** | **Hemisphere** | **Cluster size** | **T value** | **MNI X** | **MNI Y** | **MNI Z** |
| --- | --- | --- | --- | --- | --- | --- |
| Calcarine (V1) | R | 2991 | 5,70 | 20 | -94 | 2 |
| Occipital cortex mid | R |  | 5,34 | 38 | -82 | -6 |
| Lingual gyrus | R |  | 5,13 | 18 | -90 | -8 |
| Occipital cortex inf | R |  | 5,08 | 30 | -86 | -4 |
| Occipital cortex sup | L | 3013 | 5,20 | -12 | -96 | 6 |
| Occipital cortex mid | L |  | 5,12 | -30 | -76 | 24 |
| Occipital cortex mid | L |  | 5,11 | -24 | -88 | 14 |
| Parietal lobule sup | L | 77 | 3,76 | -20 | -68 | 48 |
| Parietal lobule sup | L |  | 3,22 | -22 | -62 | 42 |
| Cuneus | R | 37 | 3,69 | 16 | -80 | 42 |
| Parietal lobule sup | R | 166 | 3,67 | 26 | -72 | 52 |
| Parietal lobule sup | R |  | 3,67 | 26 | -60 | 56 |
| Parietal lobule inf | L | 36 | 3,65 | -28 | -54 | 42 |

V1: primary visual cortex; sup: superior; inf: inferior.
